# Combining Cas9 and dCas9 facilitates genome editing in genes associated with viability or welfare issues, or within paralogous gene clusters

**DOI:** 10.64898/2026.05.05.721005

**Authors:** Skevoulla Christou, Connor Macfarlane, Adam Caulder, Gemma F. Codner, Sarah N. Dowding, Matthew Mackenzie, Jade Desjardins, Karen J. Liu, Anthony R. Isles, Michelle E. Stewart, Sara Wells, Lydia Teboul

## Abstract

The high efficiency of genome editing presents a challenge when modifying genes associated with viability, welfare, or fertility issues, as implementation of the technology frequently results in mosaic animals with bi-allelic mutations. Combining deactivated Cas9 (dCas9) with Cas9 has been proposed as a strategy to protect one of the two target alleles from editing. We piloted this strategy with 11 genes that are reported as homozygous lethal or associated with welfare issues. We showed that the viability of founders was significantly increased when using 80:20 or 90:10 dCas9:Cas9 ratios, whereas the 70:30 ratio did not yield an equivalent protective effect. The associated overall production rate of mutated founder per manipulated embryo was significantly higher for the 80:20 ratio. Concomitantly, an increased proportion of dCas9 was associated with a significant increase in retention of unedited target alleles but, importantly, did not hinder germline transmission. In addition, editing genes in a paralog cluster with a combination of dCas9 and Cas9 reduced unwanted off-target editing, illustrating a further potential applicability of this approach. This study defines the optimal ratio between dCas9 and Cas9 for strategies aimed at achieving mono-allelic mutations within mosaic founders and proposes a means to reduce the incidence of off-target effects in experiments with limited gRNA options.

## Introduction

Application of CRISPR/Cas9 to genome engineering has enabled the direct introduction of targeted mutations into embryos through the simple delivery of Cas9–gRNA complexes and a homology directed repair (HDR) DNA template (1–3). Whilst this approach has significantly enhanced the efficiency of mutagenesis, high efficiency of editing can also present several challenges. Another concern is that Cas9 activity may not be strictly limited to the intended target locus. The CRISPR ribonucleoprotein (RNP) complex can show activity at genomic sites that differ from the target sequence by only a few mismatches, leading to off-target cleavage events. This makes the selection of guide RNAs (gRNAs) with minimal sequence similarity to other genomic regions a critical aspect of experimental design (4) and makes a case for the use of Cas9 variant enzymes with superior specificity (5).

In addition to off-target activity, the temporal persistence of the CRISPR/Cas9 complex within the embryo can extend beyond the window of a double-strand break (DSB) repair, increasing the risk of re-cleaving newly edited alleles (6). This reprocessing can compromise the stability of the intended sequence change. To mitigate this, blocking mutations such as alterations that disrupt the protospacer adjacent motif (PAM) corresponding to the utilised gRNA can be incorporated into the donor DNA template alongside the desired mutation, (6). These modifications prevent further recognition and cleavage of the edited allele by the residual Cas9–RNP complex.

The high efficiency of Cas9-mediated cleavage also frequently results in the simultaneous modification of both alleles at the target locus. While this may be advantageous in some contexts, it can be a significant drawback in others; for example, in cell culture systems in which a heterozygous mutation is required or when targeting genes whose complete loss of activity results in embryonic lethality or severe phenotypes (7). Notably, systematic phenotyping of knockout mice by the International Mouse Phenotyping Consortium has shown that approximately one-third of the mouse protein-coding genome is essential for viability, with homozygous mutations in these genes leading to embryonic or perinatal lethality (7, 8). In such cases, the generation of homozygous or compound heterozygous mutants in founder (G_0_) animals can hinder the recovery of viable lines (9) and produce animals with welfare issues (10). To address this, strategies have been developed to increase the likelihood of heterozygous editing outcomes within mosaic founder animals. One such approach involves co-delivering two distinct donor templates: one containing both the desired mutation and a blocking mutation, and another containing only the blocking mutation (11). This strategy aims to produce trans-heterozygous alleles, where both alleles carry the blocking mutation but only one harbours the desired edit. However, this outcome depends on both alleles being repaired via HDR, which is generally less frequent an outcome than non-homologous end joining (NHEJ) (10–12). Alternatively, adjusting the distance between the Cas9 cut site and the intended mutation has been proposed to influence the ratio of heterozygous to homozygous editing events (6). However, this approach is constrained by the availability of suitable PAM sequences near the target site. The technically challenging pronuclear transplantation method has even been used to achieve mono-allelic mutation (13), underscoring the need for simpler methods to edit genes associated with viability, welfare, or breeding issues.

Targeting a single gene within a well-conserved gene family in cell culture presents additional challenges, particularly as subsequent segregation of alleles through breeding is not possible. In such cases, paralogous genes may be unintentionally targeted due to sequence similarity, increasing the risk of unintended edits. This challenge has conventionally been addressed by increasing the number of edited clones screened to identify those with the desired mutation(14). Similarly, *in vivo* editing of one or a subset of genes within a paralogous cluster can result in off-target activity at close proximity to the target locus, complicating the generation of specific mutants. The generation of a single, specific mutation in these contexts may require multiple targeting attempts to isolate the desired allele.

Yehia and colleagues recently presented a proof of principle termed deactivated Cas9 sequestration CRISPR (das-CRISPR; TE-5 in (15) and Figure 1), designed to promote monoallelic editing through the co-delivery of catalytically active Cas9 and a nuclease-dead Cas9 variant (dCas9), in which both the RuvC and HNH nuclease domains are inactivated (16): The authors demonstrated that an 80:20 dCas9:Cas9 ratio supports monoallelic mutagenesis in proof-of-concept experiments conducted in vitro and in embryos, whereas a 50:50 ratio failed to provide the same protective effect. We piloted the use of dCas9 and Cas9 co-delivery in 11 genome-editing projects aimed at introducing single nucleotide variants (SNVs) into genes annotated as homozygous lethal or associated with welfare or fertility concerns in mice (Supplementary Table 1). We extended the range of dCas9:Cas9 molar ratios (0:100, 70:30, 80:20, and 90:10) to identify conditions that would increase the likelihood of recovering viable founders while maintaining efficient mutagenesis. We show that high dCas9:Cas9 ratios (≥80:20) improve the yield of mutated founder animals that retain wild-type alleles and enable successful generation of founder animals in projects where previous attempts had failed. Building on our success in homozygous lethal projects, we show that this approach can also be applied to reduce off-target editing within gene clusters, facilitating the specific modification of individual paralogs.

**Figure 1:**
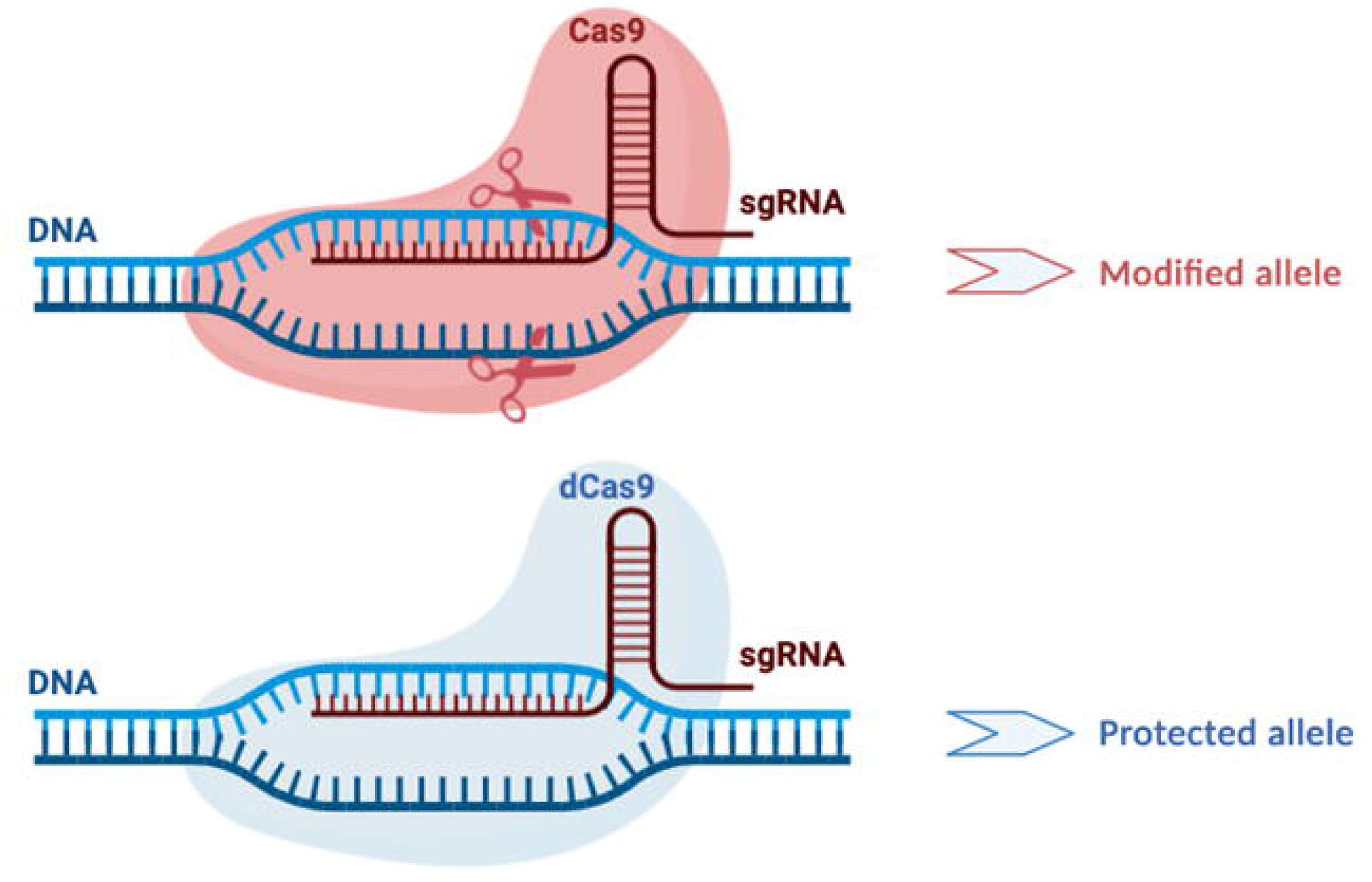
Schematic of functional wild-type Cas9 (pink, top) versus deactivated Cas9 (dCas9) (blue, bottom). The WT Cas9 will cause a double-stranded break at the site the guide RNA has bound to leading to endogenous DNA repair pathways being activated. The dCas9 will also bind at the site the guide RNA is homologous to but does not have endonuclease activity so does not cause any breaks to the DNA. This protects the allele from the effects of the WT Cas9 and will not activate DNA repair pathways. Created in BioRender.com

## Materials and method

### Design of sgRNAs and oligonucleotide templates

Guide sequence selection was carried out using the following online tools: CRISPOR (17) and WTSI Genome Editing (WGE) (18). Chemically synthesised single guide (sg)RNA (modified CRISPR RNA containing stabilising 2’-O-methyl and phosphorothioate linkages purified via reverse-phase cartridge chromatography) were obtained from Merck. Sequences for single-stranded oligo-deoxynucleotide (ssODN) HDR templates were designed with homology arms at least 60 nucleotides (nt) in length flanking the intended point mutation. These were generally centred on the CRISPR cutting site with exceptions offset towards a point mutation to favour a recombination event that includes the desired change. Wherever possible, silent point mutations were introduced to the ssODN HDR template sequence to disrupt the PAM sequence that is recognised by the CRISPR RNP and prevent re-cutting of the modified allele by Cas9. If alteration of the PAM sequence was not possible, further changes in the seed sequence within the protospacer to which the sgRNA will bind were introduced to the ssODN template. All ssODN templates were ordered as 200-mer Ultramer™ DNA oligonucleotides (IDT) with four phosphorothioate modifications placed at each of the 5′ and 3′ extremities to increase stability (12, 19). On reception, oligonucleotides were re-suspended in microinjection buffer (10 mM Tris-HCl, 0.1 mM EDTA, 100 mM NaCl, pH7.5). The sequences of oligonucleotides, protospacers and ssODN HDR templates used within the examples presented in this study are shown in Supplementary Table 2.

### Mice

All animals were housed and maintained in the Mary Lyon Centre, MRC Harwell under specific pathogen-free (SPF) conditions, in individually ventilated cages adhering to environmental conditions as outlined in the Home Office Code of Practice. All animal studies were licenced by the Home Office under the Animals (Scientific Procedures) Act 1986 Amendment Regulations 2012 (SI 4 2012/3039), UK, and additionally approved by the Institutional Ethical Review Committee. Mice, including those showing welfare issues reaching humane endpoints, were euthanised by Home Office Schedule 1 methods.

### Embryo electroporation

All embryos were obtained by superovulation and mating of C57BL/6J or C57BL/6NTac mice. Separate preparations of dCas9 (IDT):Cas9 protein (Sigma) were made in advance to the following weight ratios: 0:100, 70:30, 80:20 and 90:10 from 10 μg/μl dCas9 protein and 5 μg/μl Cas9 protein stocks. The dCas9:Cas9 combination, sgRNAs (Sigma) and ssODN HDR templates (IDT) were diluted and mixed in Electroporation buffer (EB; Gibco Opti-MEM I Reduced Serum Media – (Thermo Fisher Scientific)) to final concentrations of 650 ng/μl total, 130 ng/μl total and 400 ng/μl, respectively. Embryos were electroporated using the following conditions: 40 V, 3.5 ms pulse length, 50 ms pulse interval, 4 pulses (NEPA21 Type II – (NEPA Gene)). Mixes were centrifuged at high speed for one minute prior to use for electroporation. Electroporated embryos were transferred into CD-1 pseudopregnant females. Host females were allowed to litter and rear G_0_ animals. In all cases, mixes corresponding to at least two different dCas9:Cas9 ratios were used on a given day for each editing project.

### Genotyping

Genomic DNA from G_0_ and G_1_ animals was extracted from ear clip biopsies using the DNA Extract All Reagents Kit (Applied Biosystems) according to manufacturer’s instructions. The crude lysate was stored at -20°C. G_0_ and G_1_ animals were screened for the presence of the desired mutation, and genotyped by PCR amplification of the region of interest and Sanger sequencing as previously described (20) or with a combination of Sanger (sourced from Source Bioscience) and Oxford Nanopore Technology (sourced from Plasmidsaurus) sequencing (21), respectively. To exclude animals with additional HDR template copies, the presence of additional integrations of HDR templates was screened for using ddPCR in G_1_ animals (13). The sequences of primers and probes used are shown in Supplementary Table 2.

### Breeding for germline transmission

G_0_ animals in which the presence of a desired allele was detected were mated to wild-type (WT) isogenic animals to obtain G_1_ animals, in which to assess the germline transmission of the allele of interest and complete the validation of its integrity.

### Data analysis and availability

Data are expressed as a percentage of the number of embryos electroporated and transferred or as a percentage of the number of animals analysed. The number of corresponding electroporation sessions across the 11 projects for data expressed as a percentage of embryos electroporated and transferred were 17, 10, 21 and 12 for the 0:100, 70:30, 80:20 and 90:10 dCas9:Cas9 conditions, respectively. As some sessions resulted in no pups reaching weaning stage, the number of corresponding electroporation sessions across the 11 projects for data expressed as a percentage of the number of animals analysed were 7, 8, 20 and 11 for the 0:100, 70:30, 80:20 and 90:10 dCas9:Cas9 conditions, respectively. All graphs and statistics were produced and calculated using GraphPad Prism Software (version 10.5.0). Data were visualised as box and whisker plots whereby the median, 25^th^ and 75^th^ percentiles are displayed by the box, and the whiskers show the maximum and minimum data points. Descriptive statistics were performed and included basic parameters as well as skewness and kurtosis. Skewness (measurement of the symmetry of data distribution) was calculated using the G_1_ method. Values of more than 1 or less than –1 indicate that there was substantial asymmetrical distribution of the data. Kurtosis is used to assess whether the distribution of data in the tails is Gaussian. Positive kurtosis indicates the presence of more data in the tails than is observed in normally distributed data, whereas negative kurtosis indicates less data in the tails. Statistical analysis was performed using non-parametric Kruskal–Wallis Analysis of Variance (KW ANOVA) followed by Dunn’s pairwise comparison test. A *p* value of less than 0.05 was considered statistically significant. For pairwise comparison, all statistically significant results were compared with the 0:100 dCas9:Cas9 condition. All data are presented in the main text or in supplementary figures and tables.

## Results

### Combining dCas9 and Cas9 restores viability in founders for genome editing of genes associated with lethality or welfare issues

Genome-editing experiments targeting homozygous lethal genes are recognised as being particularly challenging due to poor birth rates and generation of founders with welfare issues that prevent germline transmission of the desired mutations (9, 10). We conducted 11 projects aimed at introducing SNVs into genes associated with viability or welfare concerns (Supplementary Table 1). We first assessed the survival of animals following genetic manipulation. Figure 2 presents the number of weaned animals per embryo electroporated with CRISPR RNP and ssODN HDR template and transferred, hereafter referred to as ‘animals per embryo’. Each data point represents a session in which approximately 75 embryos were electroporated and transferred into pseudopregnant females. Electroporation using Cas9 RNPs and ssODN templates alone yielded few or no potential founders for screening (Figure 2, condition 0:100; median: 0%; average: 5.9% animals per embryo; Table 1). The distribution of outcomes was skewed (skewness = 1.333; kurtosis = 0.7744), with 10 out of 17 sessions resulting in no animals reaching weaning age. Notably, initial manipulation of 1218 embryos with WT Cas9 alone across 11 projects only yielded 37 pups to analyse, showing the low return of animals of the Cas9 only condition for projects aimed at mutagenesis of homozygous lethal genes or those associated with welfare issues (Supplementary Table 3).

**Figure 2:**
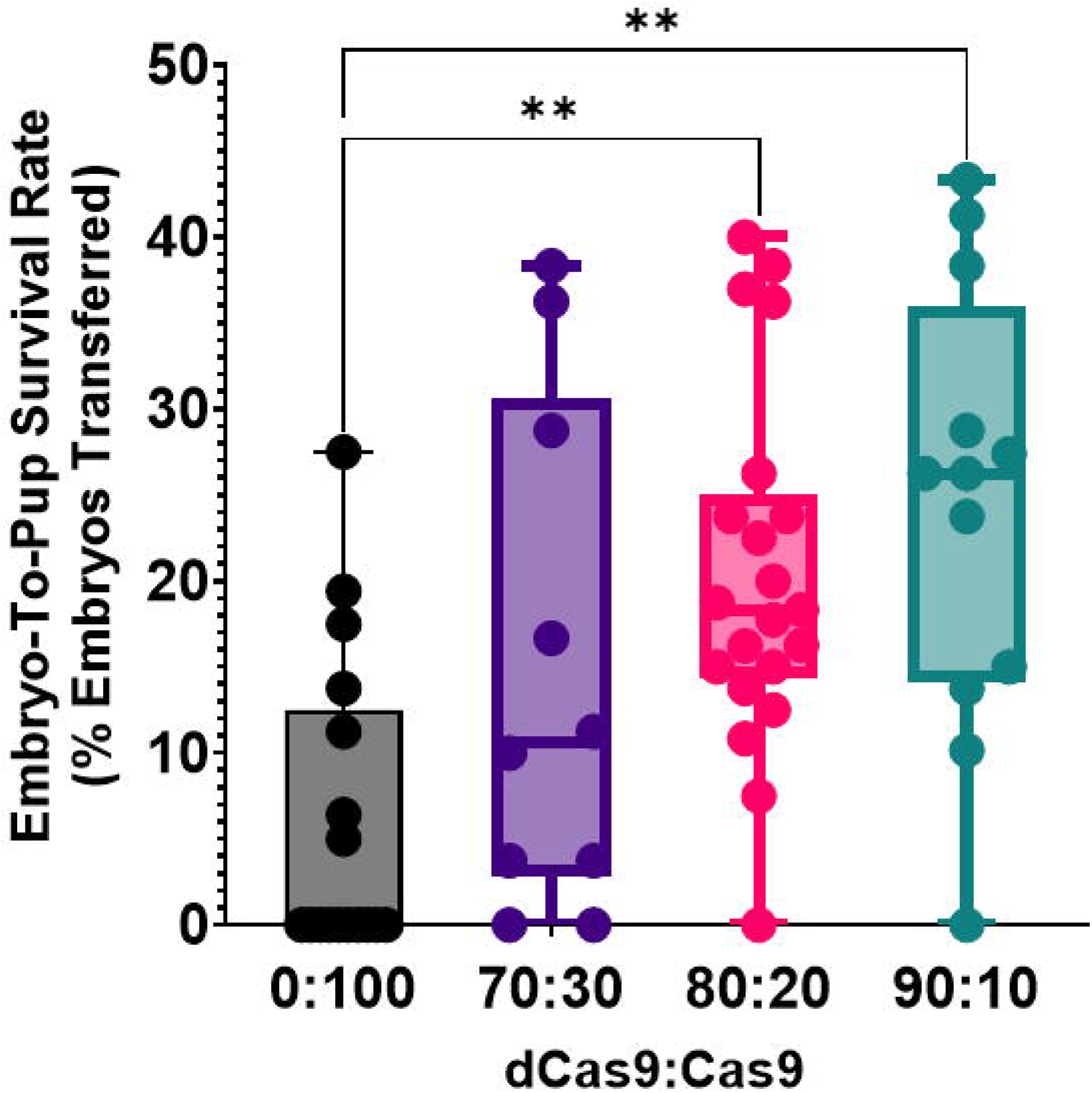
Embryo-to-pup survival rate is significantly increased in homozygous lethal genes when using increasing ratios of dCas9:Cas9. The number of pups surviving until weaning expressed as a percentage of the total number of embryos that were electroporated and transferred showed a significant difference between groups (p < 0.001). Pairwise comparison showed significant increases in embryo-to-pup survival for the 80:20 (p < 0.01), and for the 90:10 dCas9:Cas9 (p < 0.01).

**Table 1:** A summary of each dCas9:Cas9 condition showing the number of embryos used, number of pups surviving to weaning age and the number of electroporation sessions where either no pups were born, or no pups remained for analysis after weaning age. The percentage of electroporations that resulted in no pups was far higher for the WT 0:100 dCas9:Cas9 condition than for the others tested.

| **Enzyme Ratio** | **0:100** | **70:30** | **80:20** | **90:10** |
| --- | --- | --- | --- | --- |
| **No. of electroporations** | 17 | 10 | 21 | 12 |
| **No. of embryos used** | 1247 | 760 | 1609 | 932 |
| **No. of pups analysed** | 77 | 108 | 314 | 237 |
| **% pups generated from embryos electroporated** | 6.2 | 14.2 | 19.5 | 25.4 |
| **No. of electroporation sessions with zero pups born** | 6 | 2 | 1 | 0 |
| **No. of electroporation sessions with zero pups to analyse** | 10 | 2 | 1 | 1 |
| **% of electroporations that result in no pups** | 58.8 | 20 | 4.8 | 8.3 |

To address this limitation, we introduced dCas9 alongside Cas9 to increase the likelihood of generating heterozygous lineages with a functional WT allele in founder animals. This strategy leverages the ability of dCas9 to bind the target sequence and potentially shield it from binding and DSB induction by WT Cas9 (Figure 1, (22), (TE-5, 16)). We chose to assess the dCas9:Cas9 ratios 80:10 and 90:10, as proposed for embryos by others, and 70:30 to find the optimal balance between input of nuclease activity and shielding of the target locus. Substitution of Cas9 with increasing proportions of dCas9 led to a marked improvement in the recovery rate of potential founders with which to screen for the presence of the intended SNPs; a significant difference was seen between the groups (KW ANOVA, p < 0.001). Specifically, the median number of animals per embryo increased from 0% with Cas9 only to 10.6%, 18.3%, and 26.3% for 70:30, 80:20, and 90:10 dCas9:Cas9 ratios, respectively.

Pairwise comparisons revealed statistically significant increases in the yield of animals reaching weaning stage between the 0:100 (Cas9 only) and 80:20 (Dunn’s test; p < 0.01), as well as the 0:100 and 90:10 (Dunn’s test; p < 0.01) conditions (Figure 2). Although the 70:30 condition showed increased yield, it did not reach statistical significance compared to the 0:100 condition. Importantly, the distribution of outcomes in the 80:20 and 90:10 conditions was more uniform (skewness = 0.4334 and –0.2812, respectively), in contrast to the highly skewed distribution observed with Cas9 alone (skewness = 1.333). Table 1 summarises the outcomes per electroporation session and illustrates a positive correlation between increasing dCas9 proportion and the proportion of sessions yielding viable, weaned animals.

### Evaluation of mutagenesis outcome in genome-editing experiments with combinations of dCas9 and Cas9

Given the significant increase in the number of pups surviving until weaning when incorporating dCas9, we next analysed the frequency of WT alleles to evaluate the protective effect of dCas9 and the efficiency of generating founders with the desired mutations. The regions of interest were amplified by PCR from DNA extracted from G_0_ ear biopsies and was sequenced by Sanger sequencing (See Materials and methods). Genotyping outcomes were classified as WT, incorrect mutation (indels or illegitimate repair involving incorrect integration of the template sequence), or desired mutation (HDR-mediated legitimate repair) (12). G_0_ animals could show mosaicism, harbouring alleles from all three categories. The genotyping data for the 11 experiments are shown in Supplementary Figures 1 to 11.

In the absence of dCas9, the number of G_0_ animals with detectable WT alleles (either alone or in combination with mutant alleles) per embryo was low (median: 0%; Figure 3a). The inclusion of dCas9 significantly increased the number of animals with WT alleles per embryo across all tested ratios (Figure 3a; KW ANOVA, p < 0.0001; Dunn’s test vs. 0:100: 70:30, p < 0.05; 80:20 and 90:10 dCas9:Cas9, p < 0.0001). Similarly, the proportion of animals with evidence of WT alleles among those animals analysed increased significantly when using the combination of dCas9 and Cas9, in all ratios tested (Figure 3d; median: 9.1% (0:100), 98.3% (70:30), 93.1% (80:20), 100% (90:10); KW ANOVA, p < 0.0001; Dunn’s test vs. 0:100: p < 0.01 for 70:30 and 80:20; p < 0.0001 for 90:10 dCas9:Cas9).

**Figure 3:**
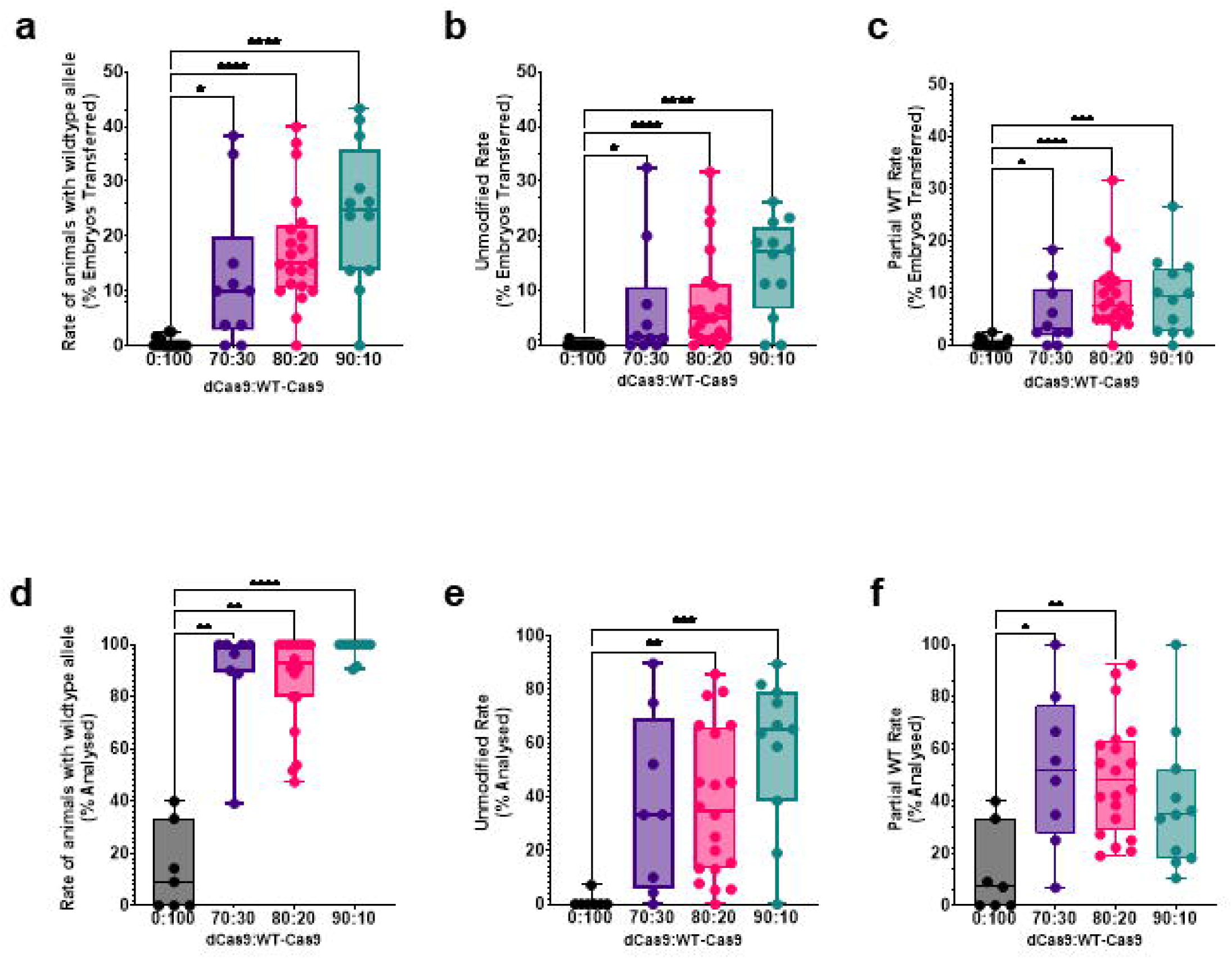
The rate of an unmodified wild-type allele significantly increases as the ratio of dCas9:Ca9 increases, highlighting the protective effect of the dCas9 from the endonuclease activity of the Cas9. (a-c) rate expressed as percentage of embryos electroporated and transferred. (d-e) rate expressed as percentage of animals analysed. (a) The rate of animals showing evidence of a wild-type (WT) allele, showed a significant difference between groups (p < 0.0001). Pairwise comparison showed significant increases in the presence of the WT allele for all the conditions tested. (b) The rate of unmodified animals where only a WT allele was detected, showed a significant difference between groups (p < 0.0001). Pairwise comparison showed significant increases in the rate of unmodified animals for all the conditions tested. (c) The rate of animals in which a WT allele was detected alongside a mutated allele, showed a significant difference between groups (p < 0.0001). Pairwise comparison showed significant increases in the rate of animals with a WT allele alongside mutated alleles for all the conditions. (d) The rate of animals showing evidence of a WT allele, showed a significant difference between groups (p < 0.0001). Pairwise comparison showed significant increases in the presence of the WT allele for all the conditions tested. (e) The rate of unmodified animals in which only a WT allele was detected, showed a significant difference between groups (p < 0.001). Pairwise comparison showed significant increases in the rate of unmodified animals for the 80:20 and 90:10 dCas9:Cas9 conditions. Pairwise comparison showed significant increases in the rate of unmodified animals for the 80:20 and 90:10 dCas9:Cas9 conditions. (f) The rate of animals in which a WT allele was detected alongside a mutated allele, showed a significant difference between groups (p < 0.01). Pairwise comparison showed significant increases in rate of animals with a WT allele alongside mutated alleles for the 70:30 and 80:20 dCas9:Cas9 conditions. Pairwise comparison showed significant increases in the rate of animals with a WT allele alongside mutated alleles for the 70:30 and 80:20 dCas9:Cas9 conditions. * p < 0.05, ** p < 0.01, *** p < 0.001 and ****p < 0.0001.

We next interrogated the efficiency of the strategy by evaluating the production of animals with no evidence of mutagenesis (only WT allele detected). The number of animals with only WT allele detected per embryo also increased significantly with the use of dCas9 (Figure 3b; KW ANOVA, p < 0.0001; Dunn’s test vs. 0:100: 70:30, p < 0.05; 80:20 and 90:10 dCas9:Cas9, p < 0.0001). As the proportion of dCas9 increased, so did the percentage of unedited founders per embryo (Figure 3b; median: 0% (0:100), 1.5% (70:30), 5% (80:20), 17.1% (90:10)). Similar proportional trends were observed for the presence of only WT allele as per animals analysed (Figure 3e; KW ANOVA, p < 0.01; Dunn’s test vs. 0:100: 80:20, p < 0.01; 90:10 dCas9:Cas9, p < 0.001).

We next assessed whether the use of dCas9, which enhanced founder viability, limited the generation of mutant alleles. To address this, we evaluated the rate of animals harbouring both WT and mutant alleles per embryo, in order to identify ratios that balance the protective effect of dCas9 with sufficient Cas9 activity. This rate increased significantly with dCas9 (Figure 3c; KW ANOVA, p < 0.0001; Dunn’s test vs. 0:100: 70:30, p < 0.05; 80:20, p < 0.0001; 90:10 dCas9:Cas9, p < 0.001). Similar significant increases were seen for the number of animals bearing both WT and mutant alleles expressed as a percentage of animals analysed for the 70:30 and 80:20 dCas9:Cas9 conditions (Figure 3f; KW ANOVA, p < 0.01; Dunn’s test vs. 0:100: 70:30, p < 0.05; 80:20 dCas9:Cas9, p < 0.01).

We next evaluated the overall mutation rate, defined as the number of founders carrying at least one mutant allele per embryo (Figure 4a) and per animal analysed (Figure 4b) highlighting the high mutagenesis efficiency of the CRISPR–Cas9 RNPs. Cas9 alone produced a median mutation rate of 0% per embryo (Figure 4a), whereas the median mutation rate per animal analysed was 100% (Figure 4b). The data were skewed per embryo and per animal analysed (skewness: 1.36 and –2.65; kurtosis: 0.93 and 7.00, respectively). The 80:20 condition showed the highest median (10%) and average (12.3%) mutation rate per embryo, with a significant increase over Cas9 alone (KW ANOVA, p < 0.05; Dunn’s test vs. 0:100, 80:20 dCas9:Cas9 p < 0.05). When expressed per animal analysed, mutation rates decreased significantly with increasing dCas9 ratio (Figure 4b; KW ANOVA, p < 0.001; Dunn’s test vs. 0:100: 80:20, p < 0.05 and 90:10 dCas9:Cas9, p < 0.001). This reflects the increased number of unmutated animals born in conditions in which dCas9 was used.

**Figure 4:**
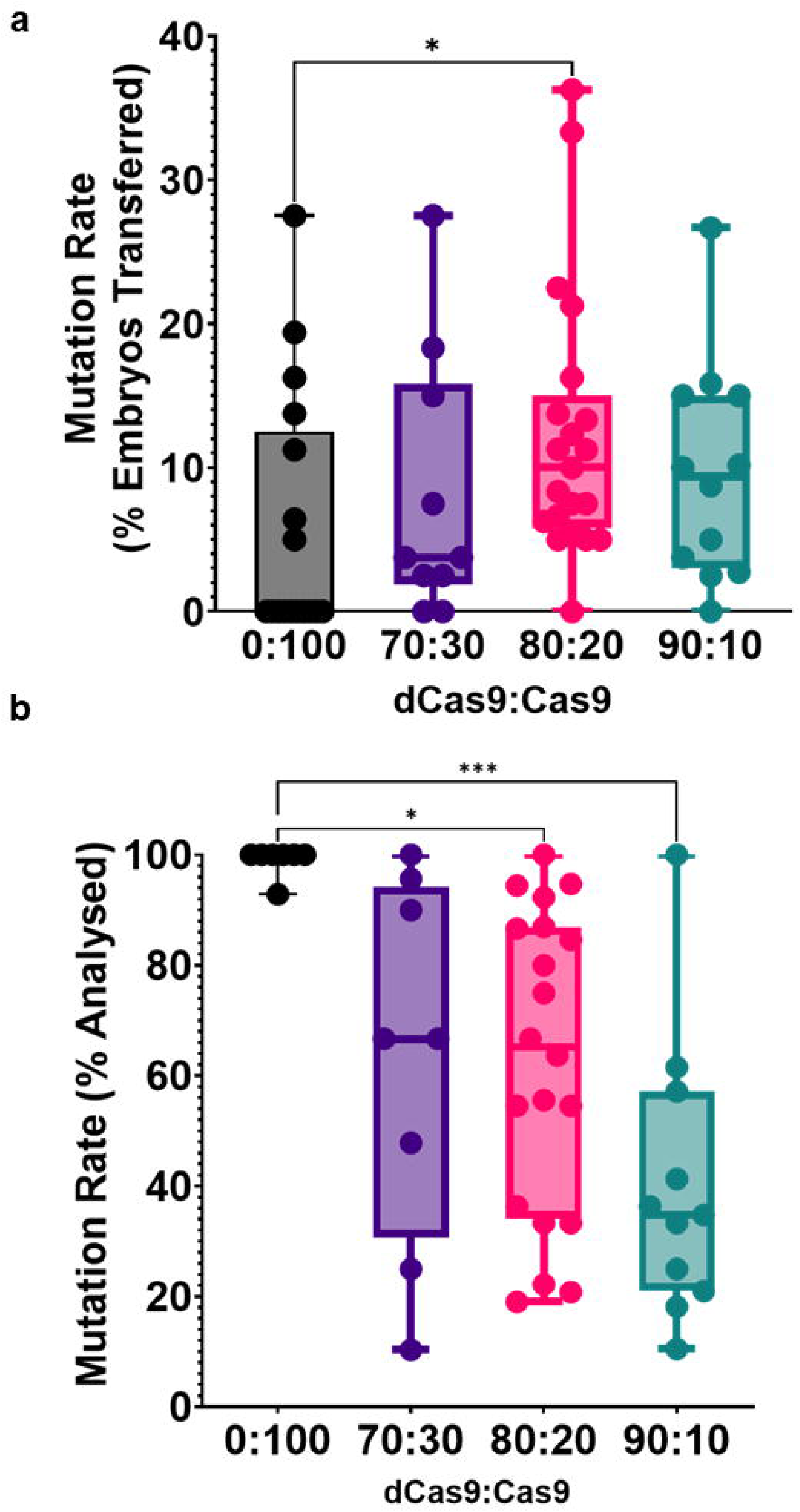
Analysis of mutations in founders. (a) The rate at which a mutant allele was seen in animals, expressed as the percentage of embryos electroporated and transferred, showed a significant difference between groups (p < 0.05). Pairwise comparison showed significant increases in the presence of mutant alleles for the 80:20 dCas9:Cas9 condition (p < 0.05). The median mutation rate was 3.8% (70:30), 10.0 % (80:20) and 9.4% (90:10) compared with 0 % for the (0:100) dCas9:Cas9 condition. The average mutation rate was 8.1 % (70:30),12.3 % (80:20) and 9.6 % (90:10) compared with 5.9 % for the (0:100) dCas9:Cas9 condition. (b) The rate at which a mutant allele was seen in animals, expressed as the percentage of animals analysed, showed a significant different between the conditions (p < 0.001). Pairwise comparison showed a significant decrease in the presence of mutant alleles for the 80:20 (p < 0.05) and 90:10 (p < 0.001) dCas9:Cas9 conditions. The median mutation rate was 66.7 % (70:30), 65.2 % (80:20) and 34.8 % (90:10) compared with 100 % for the (0:100) dCas9:Cas9 condition. The average mutation rate was 62.8 % (70:30), 62.8 % (80:20) and 39.9 % (90:10) compared with 99.0 % for the (0:100) dCas9:Cas9 condition.

Finally, we assessed the rate of desired HDR events. No significant differences were observed in overall rates of animals with HDR per embryo across conditions (Figure 5a; (median: 0% (0:100), 1.9% (70:30), 2.5% (80:20) and 1.5% (90:10)). When assessed per analysed animal, overall HDR rates were significantly lower with the 90:10 ratio compared to Cas9 only, while no significant difference was seen between the other conditions (Figure 5b; KW ANOVA, p < 0.05, Dunn’s test vs 0:100, 90:10 dCas9:Cas9, p < 0.05). The average desired HDR rate among mutant animals decreased with increasing dCas9 ratios: Cas9 only (55.6%), 70:30 (40.7%), 80:20 (26.8%), and 90:10 (20.2%) (Figure 5b). It is noteworthy that the ratio of animals with desired mutation per animal analysed could not be calculated for 14 out of 60 sessions as these did not yield any animals for analysis. Ten of these sessions corresponded to the Cas9 only condition, whereas there were 2 sessions for the 70:30, 1 for the 80:20 and 1 for the 90:10 dCas9:Cas9 conditions, respectively (Table 1). Although the average desired HDR yields per embryo did not differ significantly (average: 3.3% (0:100), 2.5% (70:30), 3.4% (80:20) and 2.3% (90:10)), the distribution of positive outcomes varied. Cas9 alone yielded positive founders in 35.3% of sessions, while 70:30, 80:20, and 90:10 dCas9:Cas9 conditions yielded 60%, 76.2%, and 58.3% of sessions with positive founders, respectively (Figure 5b, Table 2). Importantly, all 11 projects produced positive founders with the 80:20 condition, compared to only 4 with Cas9 alone (Table 2).

**Figure 5:**
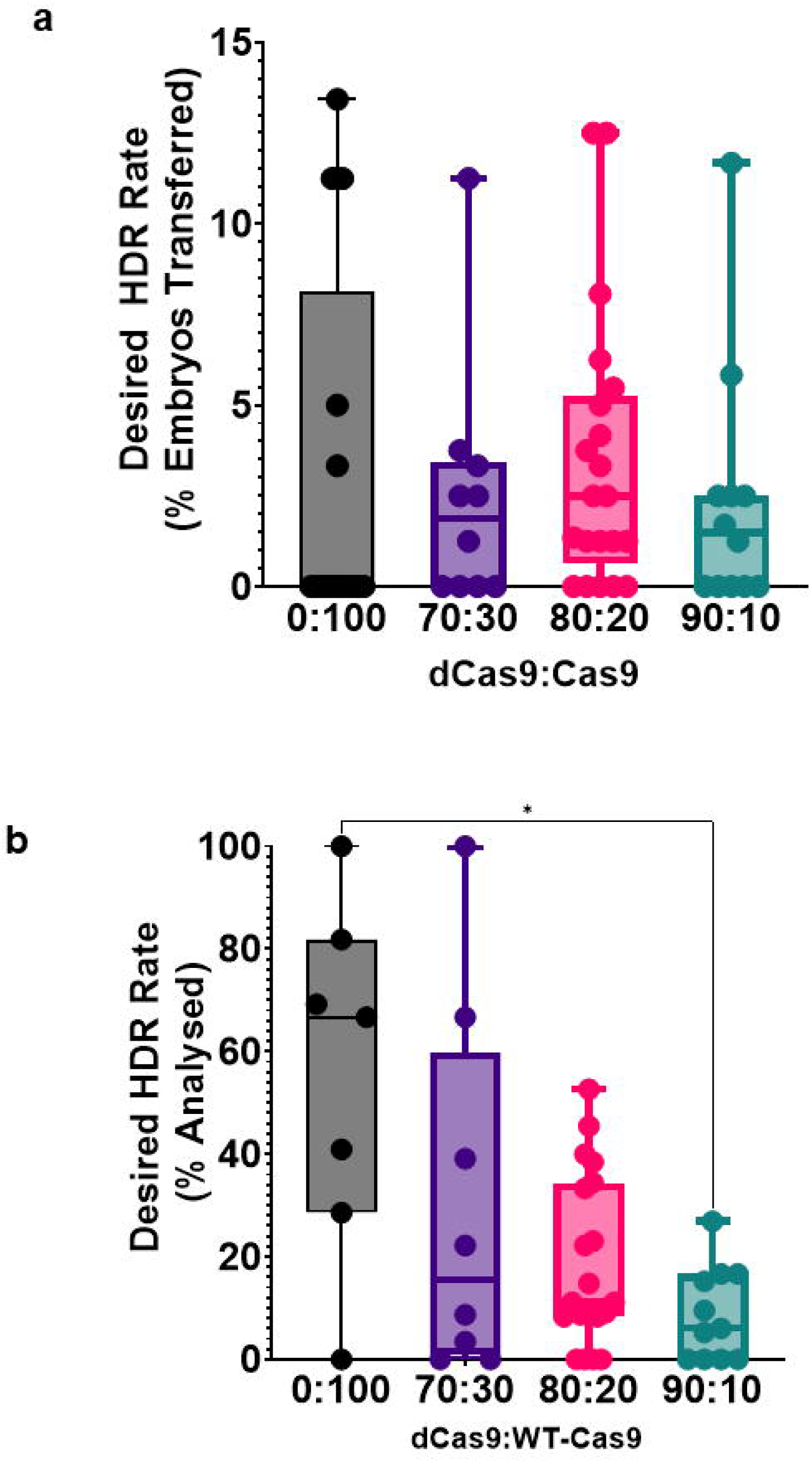
The rate of desired homology directed repair for embryos electroporated and transferred was not significantly different between the different conditions, indicating that the protective effect of dCas9 does not come at the expense of the desired mutation. (a) The rate of animals with a desired HDR allele, expressed as the percentage of embryos electroporated and transferred, did not show a significant different between the conditions (p > 0.05). (b) The rate of animals with a desired HDR allele, expressed as the percentage of animals analysed, showed a significant difference between the conditions (p < 0.05). Pairwise comparison showed a significant decrease in the rate of desired HDR alleles for the 90:10 dCas9:Cas9 condition compared with the 0:100 condition (p < 0.05).

**Table 2:** A summary of each dCas9:Cas9 condition showing the rate of desired HDR against the number of electroporations and the different projects tested. The highest rate of desired HDR was seen for the 80:20 dCas9:Cas9 ratio for which all projects tested resulted in the desired mutation.

| dCas9:<br>Cas9 | No. embryos<br>electroporated<br>and<br>transferred | No. of<br>Electroporations | No.<br>electroporations<br>with desired<br>HDR | Electroporations<br>Desired HDR<br>Rate (%) | No. of<br>Projects | No.<br>projects<br>with<br>Desired<br>HDR | Projects<br>Desired<br>HDR<br>Rate(%) |
| --- | --- | --- | --- | --- | --- | --- | --- |
| 0:100 | 1247 | 17 | 6 | 35.3 | 10 | 4 | 40 |
| 70:30 | 760 | 10 | 6 | 60 | 10 | 6 | 60 |
| 80:20 | 1609 | 21 | 16 | 76.2 | 11 | 11 | 100 |
| 90:10 | 932 | 12 | 7 | 58.3 | 10 | 6 | 60 |

### Analysis of germline transmission from founders generated with combinations of dCas9 and Cas9

To evaluate the heritability of mutations generated using combinations of dCas9 and Cas9, we assessed germline transmission (GLT) from founder animals. Due to ethical and logistical constraints, not all positive G_0_ animals were bred. Instead, the first two or three positive founders per project were selected for mating with WT animals. In total, 32 G_0_ founder animals produced from the 11 projects with 0:100, 80:20 or 90:10 dCas9:Cas9 conditions, were bred to assess transmission of the desired allele. These ratios were chosen as they corresponded to a significant improvement in the embryo-to-pup survival rate. Of these, 7 founders failed to produce viable progeny, including 3 out of 5 from the 0:100 condition, 3 out of 18 from the 80:20 condition, and 1 out of 9 from the 90:10 condition (Table 3).

**Table 3:** Summary of the genotype of G_0_ founders progeny. Average % transmission is the average rate that a founder from a particular dCas9:Cas9 condition transmitted the allele to its progeny across different projects. Noticeably, the G0 founders from the 0:100 condition did not contain a WT allele to transmit to the next generation. Note that no 70:30 founders were put into breeding. *For the 80:20 dCas9:Cas9 condition, one of the projects that did not produce progeny was the MGI ID 3584508 project as G0 founders failed to produce viable progeny and lethality was seen when bred to other background strains. The other 80:20 dCas9:Cas9 condition project that did not produce progeny was the MGI ID 1919769 as no breeding attempts were made for the founder.

| Condition | Number of mated G<sub>0</sub> founders (with evidence of WT allele) | Number of G<sub>0</sub> founders that did not produce G<sub>1</sub> progeny | Number G<sub>1</sub> progeny | Number of founders with GLT | Average % transmission of desired allele | Average % transmission of other mutant allele(s) | Average % transmission of WT allele |
| --- | --- | --- | --- | --- | --- | --- | --- |
| **0:100** | 5 (0) from 3 projects | 3 from 2 projects | 47 | 2 from 2 projects | 29.1 | 70.9 | 0.0 |
| **80:20** | 18 (15) from 10 projects | 3 from 2 projects* | 353 | 6 from 6 projects | 32.2 | 35.4 | 44.2 |
| **90:10** | 9 (9) from 5 projects | 1 from 1 project | 255 | 5 from 3 projects | 26.8 | 0.8 | 73.1 |

Table 3 summarises the number of founders mated, their GLT success, and the average transmission rates of the desired, other mutant, and WT alleles. Notably, founders from the 0:100 condition that were genotyped as lacking WT alleles did not transmit WT alleles, in keeping with the genotyping results obtained from these animals. By contrast, founders from the 80:20 and 90:10 conditions showed higher WT allele transmission rates, in keeping with the genotyping results and consistent with the protective effect of dCas9. The 90:10 condition yielded an average WT allele transmission rate of 73.1%, whereas the 80:20 condition showed a consistent transmission profile across all allele types. The data show that transmission of the desired allele could be obtained by mating positive founder animals, irrespective of the conditions under which they were produced.

Analysis of potential off-target activity was not performed for the 11 projects, as no candidate off-target sites met the laboratory’s quality criteria—defined as having 2 or fewer mismatches in the protospacer sequences.

### Hedging mutagenesis events in a cluster of paralogs

To further explore the utility of dCas9 in challenging genomic contexts, we investigated its role in editing genes within clusters of highly conserved paralogs. Editing a single gene in such clusters is difficult due to the presence of high sequence similarity, which increases the risk of off-target activity, closely linked to the target sequence. In one project, we targeted the *Ear1* and *Ear2* genes, which are part of a paralogous cluster containing additional highly conserved genes. Standard Cas9-based editing yielded multiple founders in which mutations were detected for both *Ear1* and *Ear2.* Of the 27 G_0_ founders generated for *Ear1*, where the gRNA was specific to the locus, 4 founders also had evidence of editing at the *Ear2* locus (14.8%) (Supplementary Figure 12). Three G_0_ founders were taken forward to assess for successful transmission and two contained mutations in both *Ear1* and *Ear2*. However, neither founder transmitted mutations in both genes simultaneously, suggesting that co-editing events were rare or occurred *in trans* (Supplementary Table 4). When *Ear2* was targeted, the 20 G_0_ founders showed no evidence of editing in *Ear1* (Supplementary Figure 13). However, for the two founders taken forward for breeding, off-target mutations were co-transmitted with the desired *Ear2* mutation (Supplementary Table 4), strongly suggesting that the mutations were *in cis* and highlighting the difficulty of specifically editing genes in paralogous clusters.

To test whether co-editing could be achieved *in cis*, we performed a second round of editing, targeting *Ear2* in animals already heterozygous for an *Ear1* indel or targeting *Ear1* in animals heterozygous for an *Ear2* indel (Table 4). Using Cas9 alone, we generated 7 G_0_ founders with mutations in both *Ear1* and *Ear2*, which were co-transmitted to 41.7% of progeny genotyped, indicating *cis* configuration. However, all of these founders also transmitted additional off-target mutations in other paralogs within the cluster (Table 4, Supplementary Table 5). By contrast, repeating the experiment with an 80:20 dCas9-Cas9 combination yielded a G_0_ founder that transmitted both *Ear1* and *Ear2* mutations without detectable off-target changes in other paralogs (Table 4). This result highlights the potential of dCas9 to reduce off-target activity in challenging genomic regions.

**Table 4:**
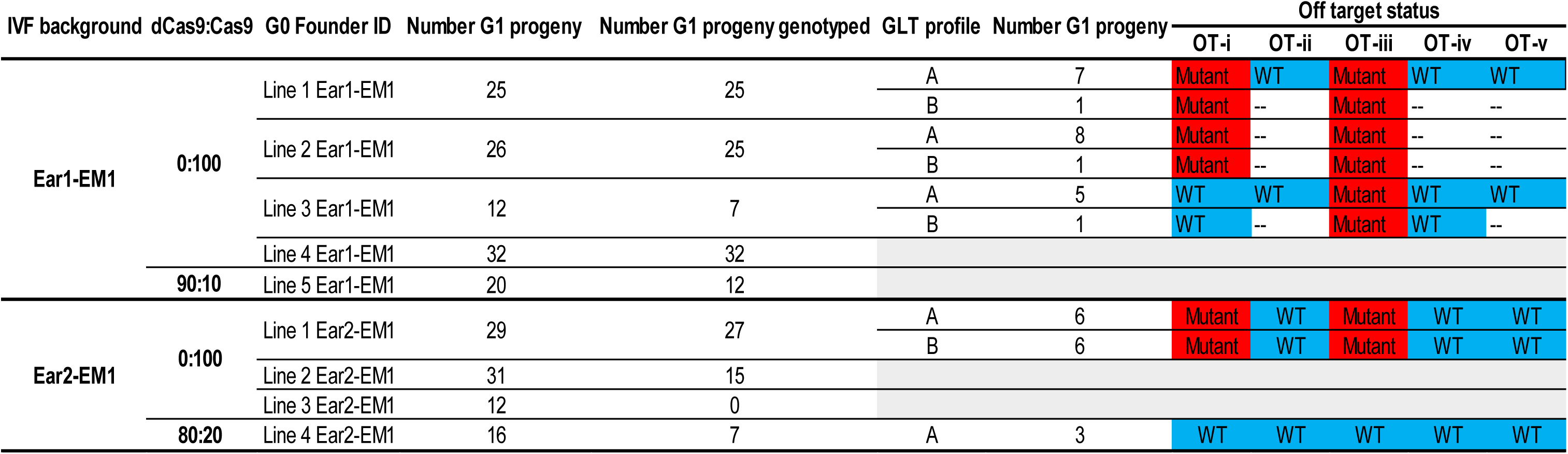
Summary of mutagenesis in a cluster of paralogs. Germ line transmission from G_0_ founders generated from the 0:100 dCas9:Cas9 condition also showed accompanying off-target mutations on the same chromosome. G_0_ founders generated from the 80:20 dCas9:Cas9 condition showed no mutagenesis at off-target sites.

## Discussion

### Efficiency of generation of viable positive founders

We performed electroporations under standard conditions for 11 editing projects, using Cas9 alone to target genes associated with viability or welfare issues. This yielded a low number of viable pups with 10 out of 17 sessions producing no pups for analysis (Figure 2, Table 1), likely due to embryonic lethality resulting from a lack of unedited alleles supporting viability. This observation is consistent with previous findings that the efficiency of generating positive founders is reduced when targeting homozygous lethal genes compared to viable ones (Supplementary Table 3 and (9)). This illustrated a significant barrier to generation of mutant mouse models. We next interrogated the potential of combining Cas9 and dCas9 to moderate the mutagenesis efficiency of Cas9 in 11 editing projects corresponding to genes associated with viability or welfare issues (Supplementary Table 1). The inclusion of increasing proportions of dCas9 in embryo electroporation significantly improved G_0_ pup yields compared to the use of Cas9 only (Figure 2, Table 1), suggesting that dCas9 competes with Cas9 at the target site, thereby reducing the likelihood of obliterating the WT allele. This approach rescued founder production in 4 of the 11 projects that had initially yielded no viable pups using Cas9 only, and increased the number of founders available for analysis in an additional 4 projects.

We next assessed whether the increased number of pups available for analysis resulted in founders with a reduced mutational burden by comparing the genotypes of animals generated using Cas9-only *versus* dCas9–Cas9 electroporation (Figure 3 and Supplementary Figures 1 to 11). While Cas9 alone yielded few founders in which a WT allele was detected, conditions incorporating dCas9 resulted in a higher proportion of founders carrying edited alleles, both correctly and incorrectly, alongside a functional WT allele (Figure 3c and 3f). These results are consistent with the proposed reduction of biallelic editing.

This reduction in mutational burden in founders would only be advantageous if the strategy retained the ability to generate mutant founders of interest. We therefore examined the overall yield of mutated animals and observed that the number of mutation-bearing founders produced per embryo electroporated was significantly higher in the 80:20 dCas9:Cas9 condition compared Cas9 alone (Figure 4a) indicating a more efficient use of embryos. Although this did not reach statistical significance, increased mutation rates were observed for the 70:30 and 90:10 dCas9:Cas9 conditions (Figure 4a), alongside an increase in viable pups. Concomitantly, mutation rates expressed as a percentage of animals analysed were lower across all dCas9:Cas9 ratios compared with Cas9 only, indicating reduced Cas9 editing activity (Figure 4b). Importantly, the distribution of mutated animals among analysed pups differed between conditions: Cas9 alone yielded consistently high mutagenesis rates of between 92% and 100%, whereas dCas9:Cas9 combinations exhibited a wider range, from 18.2% to 100%.

Despite a reduction in overall mutation rate among analysed G_0_s, with more WT animals observed (Figure 4b), the average yield of desired mutant animals per electroporated and transferred embryo did not differ significantly between ratios (Figure 5a).

However, the distribution of editing outcomes among sessions was markedly different between conditions in which different proportions of dCas9 were included. In the absence of dCas9, a high proportion of electroporation sessions yielded no viable founders for screening (Table 1), the mutagenesis rate among weaned animals was very high (Figure 4b), and fewer projects yielded positive outcomes (Table 2). By contrast, conditions that included dCas9 produced a wider range of outcomes, with a greater proportion of sessions yielding both mutants and correctly edited alleles (Table 1 and 2 and Figure 4 and 5).

Founders carrying the desired mutation were obtained in all 11 projects using the 80:20 dCas9:Cas9 ratio, compared to only 4 of 11 projects with Cas9 alone (Table 2). Importantly, founders generated in the presence of dCas9 were able to transmit the desired allele to their offspring (Table 3). These results demonstrate that the use of dCas9 represents a promising approach to support the generation of positive founders. By contrast, increasing the number of embryos used in electroporations to compensate for low founder production rates is neither practical nor ethically sustainable

### A simple solution to a common problem

Approximately 25% of protein-coding genes in the mouse genome are homozygous lethal (8), with an additional 10% associated with subviability in homozygous mutants (7). However, the introduction of SNVs is necessary to investigate protein function and validate variants of uncertain significance identified in human genetic studies (for example as discussed in (24)). As many such variants involve genes linked to viability or welfare issues, achieving viable and fertile founder animals when targeting these genes remains a frequent obstacle (10).

The combination of dCas9 and Cas9 offers a simple and effective solution for the production of these mutants. In our study, the 80:20 dCas9:Cas9 ratio supported the generation of positive founders in 16 out of 21 sessions across all 11 projects (Table 2), whereas the 90:10 ratio was effective in only a subset of cases. To account for the possibility of selecting a suboptimal sgRNA with low activity—chosen based on proximity to the desired base change—our laboratory now routinely implements a dual-electroporation strategy for projects targeting known viability- or welfare-associated genes. In this approach, the embryos are split between electroporation with Cas9 alone, and with a combination of 80:20 dCas9:Cas9.

### Effective dCas9/Cas9 ratio

An initial report using dCas9–Cas9 combinations to promote heterozygous editing used a 50:50 ratio in cultured hiPSCs (22). In the context of embryo electroporation with RNPs, another study (TE-5 in (15)), together with our data, demonstrates that higher dCas9:Cas9 ratios (80:20 or 90:10) are more effective at supporting the generation of viable, correctly edited founders (Table 1 and Table 2). This difference in effective enzyme ratios may reflect variations in delivery methods and cellular context. In embryos, both Cas9 and dCas9 are efficiently delivered via electroporation (25) and may persist, dynamically binding to and dissociating from DNA. This dynamic occupancy suggests that both alleles are likely to be exposed to active Cas9 at some point, unless dCas9 is present in sufficient excess to outcompete Cas9 for target binding. Differences in the mode of delivery, DNA repair pathways and molecular kinetics between hiPSCs and embryos may also contribute to the observed variation in optimal ratios. We also sought to optimise the strategy by testing whether a combination with incrementally more active Cas9 would show more activity whilst preserving founder viability. We found that the 70:30 dCas9:Cas9 ratio was less effective at increasing viable founder yield (Figure 2), while the 90:10 ratio resulted in fewer mutant animals as a proportion of animals analysed, than the 80:20 ratio on average (Figures 4 and 5), supporting 80:20 as an optimal default ratio. Other Cas enzymes (such as Cas12a), which exhibit different DNA binding and dissociation dynamics, may require distinct ratios between inactive and active forms.

One disadvantage of using dCas9–Cas9 combinations is the need to house and screen a larger number of founders for the desired mutation, which increases costs associated with animal maintenance, genotyping reagents and labour. However, this additional cost is justifiable when compared with the expense and delay of performing additional electroporation sessions if initial attempts using Cas9 alone fail to yield founders for screening.

### The diverse strategies to circumvent lethality and welfare issues in founders

Existing strategies to optimise genome-editing outcomes include considered gRNA design to minimise the likelihood of off-target mutations (4, 26), the use of high-specificity Cas9 variants to avoid mutation at off-target sites (5), and the application of selective DNA repair pathway inhibitors to promote HDR-mediated repair (27). However, it is reported that this latter approach may lead to additional unintended genomic alterations (28). Importantly, these strategies do not address the challenges associated with the generation of founders carrying trans-heterozygous or homozygous on-target mutations. Previous strategies to mitigate possible founder lethality have included the co-delivery of donor templates—one containing both the desired mutation and a blocking mutation, and another containing only the blocking mutation (11). This approach relies on both alleles undergoing HDR to generate a functional blocking allele and a mutant allele in a sufficient proportion of mosaic lineages. While this method allows for the generation of a useful control allele containing only the blocking mutation, it is limited by the low frequency of HDR events. By contrast, the dCas9–Cas9 strategy requires only one allele to undergo HDR, representing a more likely outcome. However, unlike the approach described by Lackner et al. (11), it does not inherently produce animals carrying only a silent mutation that can serve as controls. Finally, although pronuclear transplantation has been employed (13) to generate a mutation in the *Anapc2* gene required for preimplantation embryo development, the technical challenges associated with this approach make it unlikely to become a widely adopted strategy for targeting genes associated with viability or welfare issues.

### Hedging mutagenesis between homologous regions

We have shown that the dCas9–Cas9 approach extends beyond targeting genes associated with viability or welfare concerns. When targeting a gene within a paralog cluster, it is often difficult to design highly specific sgRNAs due to sequence similarity amongst genes and pseudogenes in the region. Here, we present an example in which a repeated *in cis* off-target effect prevented the combination of two indels in genes of the *Ear* family (Supplementary Table 4). This issue was resolved by combining Cas9 and dCas9 to retarget the locus and introduce a second *in cis* indel (Table 4), resulting in the successful addition of a single new indel at that site. This suggests that combining Cas9 and dCas9 may also lower the incidence of off-target mutations in cases where gRNAs with potential off-target risks must be used due to design constraints (Table 4).

### Improved use of animals

Genome-editing experiments targeting genes associated with viability or welfare may require large numbers of embryos to increase the probability of obtaining a favourable distribution of editing outcomes, as founder generation efficiency for these genes is generally low (Supplementary Table 3 and (9)). Here, we show that combining dCas9 and Cas9 when targeting genes associated with viability or welfare increases the yield of founders available for screening per embryo manipulated (Table1 and Figure 2), thereby reducing animal use and the number of surgeries required to generate positive founders. Using a combination of dCas9 with Cas9 results in an increase in the number of potential founders to screen for the desired mutation. However, we believe this is justifiable, as animals generated using this approach are likely to experience fewer welfare issues than those generated with Cas9 alone. Similarly, the increased production of WT G_1_ animals associated with the dCas9:Cas9 combination represents an acceptable trade-off compared to the use of animals in unproductive electroporations or the generation of trans-heterozygous animals with a high likelihood of welfare issues.

In addition, generating specific edits in a single gene within a paralogous cluster of genes often requires the production of many founder animals to identify one in which only the target of interest has been altered in the desired manner. Combing dCas9 with Cas9 reduces the proportion of animals where other genes are modified in addition to the desired target.

Overall, our data demonstrate that combining dCas9 with Cas9 is a productive strategy that reduces both mutational burden and the number of embryos required to generate founders carrying the desired mutation. This approach aligns with two of the principles of the 3Rs—Reduction and Refinement—by increasing the yield of viable, informative animals and improving the efficiency of founder generation (29).

### Relevance to biomedical research

The dCas9–Cas9 strategy is applicable to both *in vitro* (22) and *in vivo* systems, including those described in this study, and will facilitate the generation of many mutant alleles relevant to biomedical research. However, it does not address the challenge of modelling deleterious dominant-negative mutations, which may require more complex strategies such as conditional point mutations (30, 31) or conditional deletions combined with conditional overexpression alleles. The models generated in this study are currently undergoing phenotypic analysis.

## Conclusion

Combining dCas9 and Cas9 for the generation of SNVs represents a simple and effective strategy to facilitate genome editing of genes associated with lethality or welfare concerns in homozygous mutants. This approach increases the representation of WT alleles in founder animals, thereby improving the likelihood of recovering viable individuals with the desired genetic alterations. Our findings indicate that high dCas9:Cas9 ratios, specifically 80:20 or 90:10, are required under our experimental conditions to enhance the frequency of mutation-positive founder generation per manipulated embryo. Importantly, this strategy is in keeping with the principles of the 3Rs in animal research by increasing the efficiency of genome editing (Reduction) and decreasing the mutational burden in founder animals, thereby improving their welfare and the overall success rate of genome-editing experiments (Refinement) (29).

## Acknowledgements

The authors would like to thank the staff of the Mary Lyon Centre for providing excellent animal husbandry, genotyping, and microinjection services, and Dr Louise Tinsley for expert assistance with the preparation of this manuscript and Mr Shane Cody for expert graphic artwork.

---

Supplementary Table 1: Project information for genes employed in this study. Annotations sourced from MGI (https://www.informatics.jax.org/), Ensembl (https://www.ensembl.org/) and MARRVEL (https://marrvel.org/) genome browsers.

Supplementary Table 2: The sequences of sgRNAs, ssODN HDR templates, primers and probes employed in this study.

Supplementary Table 3: Summary of 0:100 dCas9:Cas9 conditions where it was the only condition tested for that day and is not included in the main analysis.

Supplementary Table 4: Summary of editing outcomes when targeting individual paralog genes, *Ear1* and *Ear2*. G_0_ founders exhibited evidence of correctly mutated alleles at both paralogs even when targeting only one site with a site-specific gRNA. However, these were not transmitted simultaneously to the G_1_ generation.

Supplementary Table 5: Off target locations for gRNAs used when targeting the *Ear1* or *Ear2* paralog genes. Off target are common to both *Ear1* and *Ear2*. Coordinates from Ensembl release 115 - September 2025.

Supplementary Figure 1: Analysis of the electroporation sessions for the MGI 3646700 point mutation project.

The figure shows the PCR amplification of the genomic region of interest with Geno_3646700_F1 and Geno_3646700_R1 (WT and mutant yields 1220 bp amplicon) for (a) session 1 with the 80:20 dCas9:Cas9 ratio, (b) session 1 with the 90:10 dCas9:Cas9 ratio, (c) session 2 with the 80:20 dCas9:Cas9 ratio, (d) session 2 with the 90:10 dCas9:Cas9 ratio and (e) session 3 with the 70:30 dCas9:Cas9 ratio from biopsies taken from the G0 animals. L1 = 1 kb DNA molecular weight ladder (thick band is 3 kb) and L2 = 100 bp DNA molecular weight ladder (thick band is 1kb). All amplicons were sent for Sanger sequencing and representative sequencing traces from (f) animal 9 from (a), (g) animal 3 from (b), (h) animal 2 from (d) and (i) animal 3 from (e) with a legitimate repair of the donor template are shown. The positions of the intended point mutations are highlighted in blue. (j) The table details the analysis of each electroporation and subsequent G0 animals with a session indicating embryos that were electroporated on the same day. The alleles seen in each pup were put into the groupings of: unmodified animals indicating no mutation event, partial wild type where the wild-type allele was seen in the presence of a mutated allele, incorrect allele where a mutated allele was present that was not desired, and desired allele repair where the repair was completed with the donor template provided. * denotes animals with evidence of allele type.

Supplementary Figure 2: Analysis of the electroporation sessions for the MGI 1917550 point mutation project.

The figure shows the PCR amplification of the genomic region of interest with Geno_1917550_F1 and Geno_1917550_R1 (WT and mutant yields 2595 bp amplicon) for (a) session 1 with the 0:100 dCas9:Cas9 ratio, (b) session 1 with the 80:20 dCas9:Cas9 ratio, (c) session 1 with the 90:10 dCas9:Cas9 ratio, (d) session 2 with the 0:100 dCas9:Cas9 ratio, (e) session 2 with the 70:30 dCas9:Cas9 ratio, and (f) session 2 with the 80:20 dCas9:Cas9 ratio from biopsies taken from the G0 animals. L1 = 1 kb DNA molecular weight ladder (thick band is 3 kb) and L2 = 100 bp DNA molecular weight ladder (thick band is 1kb). All amplicons were sent for Sanger sequencing and representative sequencing traces from (g) animal 1 from (a), (h) animal 4 from (b), (i) animal 9 from (c), (j) animal 5 from (d), (k) animal 4 from (e) and, (l) animal 2 from (f) with a legitimate repair of the donor template are shown. The positions of the intended point mutations are highlighted in blue. (m) The table details the analysis of each electroporation and subsequent G0 animals with a session indicating embryos that were electroporated on the same day. The alleles seen in each pup were put into the groupings of: unmodified animals indicating no mutation event, partial wild type where the wild-type allele was seen in the presence of a mutated allele, incorrect allele where a mutated allele was present that was not desired, and legitimate repair where the repair was completed with the donor template provided. * denotes animals with evidence of allele type. † denotes there was one less animal analysed than survived to weaning as the animal was culled prior to an ear biopsy being taken.

Supplementary Figure 3: Analysis of the electroporation sessions for the MGI 95820 point mutation project.

The figure shows the PCR amplification of the genomic region of interest with Geno_95820_F1 and Geno_95820_R1 (WT and mutant yields 969 bp amplicon) for (a) session 1 with the 0:100 dCas9:Cas9 ratio, (b) session 1 with the 80:20 dCas9:Cas9 ratio, (c) session 1 with the 90:10 dCas9:Cas9 ratio, (d) session 2 with the 70:30 dCas9:Cas9 ratio and, (e) session 2 with the 80:20 dCas9:Cas9 ratio from biopsies taken from the G0 animals. L1 = 1 kb DNA molecular weight ladder (thick band is 3 kb) and L2 = 100 bp DNA molecular weight ladder (thick band is 1kb). All amplicons were sent for Sanger sequencing and representative sequencing traces from (f) animal 12 from (a), (g) animal 16 from (b) and, (h) animal 13 from (d) with a legitimate repair of the donor template are shown. The positions of the intended point mutations are highlighted in blue. (i) The table details the analysis of each electroporation and subsequent G0 animals with a session indicating embryos that were electroporated on the same day. The alleles seen in each pup were put into the groupings of: unmodified animals indicating no mutation event, partial wild type where the wild-type allele was seen in the presence of a mutated allele, incorrect allele where a mutated allele was present that was not desired, and legitimate repair where the repair was completed with the donor template provided. * denotes animals with evidence of allele type. † denotes there was one less animal analysed than survived to weaning as the animal was culled prior to an ear biopsy being taken.

Supplementary Figure 4: Analysis of the electroporation sessions for the MGI 3584508 point mutation project.

The figure shows the PCR amplification of the genomic region of interest with Geno_3584508_F1 and Geno_3584508_R1 (WT and mutant yields 1256 bp amplicon) for (a) session 1 with the 70:30 dCas9:Cas9 ratio, (b) session 1 with the 80:20 dCas9:Cas9 ratio, (c) session 1 with the 90:10 dCas9:Cas9 ratio, (d) session 2 with the 80:20 dCas9:Cas9 ratio and (e) session 2 with the 90:10 dCas9:Cas9 ratio from biopsies taken from the G0 animals. L1 = 1 kb DNA molecular weight ladder (thick band is 3 kb) and L2 = 100 bp DNA molecular weight ladder (thick band is 1kb). All amplicons were sent for Sanger sequencing and representative sequencing traces from (f) animal 5 from (a), (g) animal 10 from (b) and, (h) animal 1 from (d) with a legitimate repair of the donor template are shown. The positions of the intended point mutations are highlighted in blue. (i) The table details the analysis of each electroporation and subsequent G0 animals with a session indicating embryos that were electroporated on the same day. The alleles seen in each pup were put into the groupings of: unmodified animals indicating no mutation event, partial wild type where the wild-type allele was seen in the presence of a mutated allele, incorrect allele where a mutated allele was present that was not desired, and legitimate repair where the repair was completed with the donor template provided. * denotes animals with evidence of allele type. † denotes there was one less animal analysed than survived to weaning as the animal was culled prior to an ear biopsy being taken. Note that for this project, a Cas9 only (0:100) condition was run on its own in a previous session and was not included in this analysis.

Supplementary Figure 5: Analysis of the electroporation sessions for the MGI 1352452 point mutation project.

The figure shows the PCR amplification of the genomic region of interest with Geno_1352452_F1 and Geno_1352452_R1 (WT and mutant yields 1155 bp amplicon) for (a) session 1 with the 80:20 dCas9:Cas9 ratio, (b) session 1 with the 90:10 dCas9:Cas9 ratio and (c) session 2 with the 80:20 dCas9:Cas9 ratio from biopsies taken from the G0 animals. L1 = 1 kb DNA molecular weight ladder (thick band is 3 kb) and L2 = 100 bp DNA molecular weight ladder (thick band is 1kb). All amplicons were sent for Sanger sequencing and representative sequencing traces from (d) animal 16 from (b) and, (e) animal 7 from (c) with a legitimate repair of the donor template are shown. The positions of the intended point mutations are highlighted in blue. (f) The table details the analysis of each electroporation and subsequent G0 animals with a session indicating embryos that were electroporated on the same day. The alleles seen in each pup were put into the groupings of: unmodified animals indicating no mutation event, partial wild type where the wild-type allele was seen in the presence of a mutated allele, incorrect allele where a mutated allele was present that was not desired, and legitimate repair where the repair was completed with the donor template provided. * denotes animals with evidence of allele type.

Supplementary Figure 6: Analysis of the electroporation sessions for the MGI 2448501 point mutation project.

The figure shows the PCR amplification of the genomic region of interest with Geno_2448501_F1 and Geno_2448501_R1 (WT and mutant yields 1066 bp amplicon) for (a) session 1 with the 0:100 dCas9:Cas9 ratio, (b) session 1 with the 80:20 dCas9:Cas9 ratio, (c) session 1 with the 90:10 dCas9:Cas9 ratio, (d) session 2 with the 0:100 dCas9:Cas9 ratio, (e) session 2 with the 70:30 dCas9:Cas9 ratio and (f) session 2 with the 80:20 dCas9:Cas9 ratio from biopsies taken from the G0 animals. L1 = 1 kb DNA molecular weight ladder (thick band is 3 kb) and L2 = 100 bp DNA molecular weight ladder (thick band is 1kb). All amplicons were sent for Sanger sequencing and representative sequencing traces from (g) animal 7 from (a), (h) animal 7 from (b), (i) animal 15 from (c), (j) animal 2 from (d), (k) animal 2 from (e) and, (l) animal 13 from (f) with a legitimate repair of the donor template are shown. The positions of the intended point mutations are highlighted in blue. (m) The table details the analysis of each electroporation and subsequent G0 animals with a session indicating embryos that were electroporated on the same day. The alleles seen in each pup were put into the groupings of: unmodified animals indicating no mutation event, partial wild type where the wild-type allele was seen in the presence of a mutated allele, incorrect allele where a mutated allele was present that was not desired, and legitimate repair where the repair was completed with the donor template provided. * denotes animals with evidence of allele type.

Supplementary Figure 7: Analysis of the electroporation sessions for the MGI 1919769 point mutation project.

The figure shows the PCR amplification of the genomic region of interest with Geno_1919769_F1 and Geno_1919769_R1 (WT and mutant yields 1007 bp amplicon) for (a) session 1 with the 80:20 dCas9:Cas9 ratio and (b) session 1 with the 90:10 dCas9:Cas9 ratio from biopsies taken from the G0 animals. L1 = 1 kb DNA molecular weight ladder (thick band is 3 kb) and L2 = 100 bp DNA molecular weight ladder (thick band is 1kb). All amplicons were sent for Sanger sequencing and representative sequencing traces from (c) animal 2 from (a), and (d) animal 17 from (b) with a legitimate repair of the donor template are shown. The positions of the intended point mutations are highlighted in blue. (e) The table details the analysis of each electroporation and subsequent G0 animals with a session indicating embryos that were electroporated on the same day. The alleles seen in each pup were put into the groupings of: unmodified animals indicating no mutation event, partial wild type where the wild-type allele was seen in the presence of a mutated allele, incorrect allele where a mutated allele was present that was not desired, and legitimate repair where the repair was completed with the donor template provided. * denotes animals with evidence of allele type.

Supplementary Figure 8: Analysis of the electroporation sessions for the MGI 105047 point mutation project.

The figure shows the PCR amplification of the genomic region of interest with Geno_105047_F1 and Geno_105047_R1 (WT and mutant yields 1740 bp amplicon) for (a) session 1 with the 80:20 dCas9:Cas9 ratio, (b) session 1 with the 90:10 dCas9:Cas9 ratio, (c) session 2 with the 70:30 dCas9:Cas9 ratio and (d) session 2 with the 80:20 dCas9:Cas9 ratio from biopsies taken from the G0 animals. L1 = 1 kb DNA molecular weight ladder (thick band is 3 kb) and L2 = 100 bp DNA molecular weight ladder (thick band is 1kb). All amplicons were sent for Sanger sequencing and representative sequencing traces from (e) animal 13 from (a), (f) animal 11 from (b), (g) animal 2 from (c) and (h) animal 15 from (d) with a legitimate repair of the donor template are shown. The positions of the intended point mutations are highlighted in blue. (i) The table details the analysis of each electroporation and subsequent G0 animals with a session indicating embryos that were electroporated on the same day. The alleles seen in each pup were put into the groupings of: unmodified animals indicating no mutation event, partial wild type where the wild-type allele was seen in the presence of a mutated allele, incorrect allele where a mutated allele was present that was not desired, and legitimate repair where the repair was completed with the donor template provided. * denotes animals with evidence of allele type.

Supplementary Figure 9: Analysis of the electroporation sessions for the MGI 1346006 point mutation project.

The figure shows the PCR amplification of the genomic region of interest with Geno_1346006_F1 and Geno_1346006_R1 (WT and mutant yields 1060 bp amplicon) for (a) session 1 with the 0:100 dCas9:Cas9 ratio, (b) session 1 with the 80:20 dCas9:Cas9 and (c) session 2 with the 80:20 dCas9:Cas9 ratio from biopsies taken from the G0 animals. L1 = 1 kb DNA molecular weight ladder (thick band is 3 kb) and L2 = 100 bp DNA molecular weight ladder (thick band is 1kb). All amplicons were sent for Sanger sequencing and representative sequencing traces from (d) animal 2 from (a), and (e) animal 12 from (c) with a legitimate repair of the donor template are shown. The positions of the intended point mutations are highlighted in blue. (f) The table details the analysis of each electroporation and subsequent G0 animals with a session indicating embryos that were electroporated on the same day. The alleles seen in each pup were put into the groupings of: unmodified animals indicating no mutation event, partial wild type where the wild-type allele was seen in the presence of a mutated allele, incorrect allele where a mutated allele was present that was not desired, and legitimate repair where the repair was completed with the donor template provided. * denotes animals with evidence of allele type.

Supplementary Figure 10: Analysis of the electroporation sessions for the MGI 1315202 point mutation project.

The figure shows the PCR amplification of the genomic region of interest with Geno_1315202_F1 and Geno_1315202_R1 (WT and mutant yields 944 bp amplicon) for (a) session 1 with the 0:100 dCas9:Cas9 ratio, (b) session 1 with the 70:30 dCas9:Cas9 ratio and (c) session 1 with the 80:20 dCas9: Cas9 ratio from biopsies taken from the G0 animals. L1 = 1 kb DNA molecular weight ladder (thick band is 3 kb) and L2 = 100 bp DNA molecular weight ladder (thick band is 1kb). All amplicons were sent for Sanger sequencing and representative a sequencing trace (d) from animal 6 from (c) with a legitimate repair of the donor template are shown. The positions of the intended point mutations are highlighted in blue. (e) The table details the analysis of each electroporation and subsequent G0 animals with a session indicating embryos that were electroporated on the same day. The alleles seen in each pup were put into the groupings of: unmodified animals indicating no mutation event, partial wild type where the wild-type allele was seen in the presence of a mutated allele, incorrect allele where a mutated allele was present that was not desired, and legitimate repair where the repair was completed with the donor template provided. * denotes animals with evidence of allele type.

Supplementary Figure 11: Analysis of the electroporation sessions for the MGI 107595 point mutation project.

The figure shows the PCR amplification of the genomic region of interest with Geno_107595_F1 and Geno_107595_R1 (WT and mutant yields 1512 bp amplicon) for (a) session 1 with the 80:20 dCas9:WT Cas9 ratio, (b) session 1 with the 90:10 dCas9:WT Cas9 ratio, (c) session 2 with the 70:30 dCas9:WT Cas9 ratio and (d) session 2 with the 80:20 dCas9:WT Cas9 ratio from biopsies taken from the G0 animals. L1 = 1 kb DNA molecular weight ladder (thick band is 3 kb) and L2 = 100 bp DNA molecular weight ladder (thick band is 1kb). All amplicons were sent for Sanger sequencing and representative sequencing traces from (e) animal 2 from (a) with a legitimate repair of the donor template are shown. The positions of the intended point mutations are highlighted in blue. (f) The table details the analysis of each electroporation and subsequent G0 animals with a session indicating embryos that were electroporated on the same day. The alleles seen in each pup were put into the groupings of: unmodified animals indicating no mutation event, partial wild type where the wild-type allele was seen in the presence of a mutated allele, incorrect allele where a mutated allele was present that was not desired, and legitimate repair where the repair was completed with the donor template provided. * denotes animals with evidence of allele type. † denotes there was one less animal analysed than survived to weaning as the animal was culled prior to an ear biopsy being taken. ‡ denotes that the animal selected for breeding had the desired point mutation but did not have the blocking mutation, indicating that it was an incorrect allele due to the repair being an incomplete use of the donor template applied.

Supplementary Figure 12: Analysis of the electroporation sessions for the *Ear1* project.

(a)The figure shows the PCR amplification of the *Ear1* genomic region of interest with Geno_Ear1_Indel_F1 and Geno_Ear1_Indel_R1 (WT yields 1842 bp amplicon) for session 1 with the 0:100 dCas9:Cas9 ratio and (b) the PCR amplification of the *Ear2* paralog genomic region of interest with Geno_Ear2_Indel_F1 and Geno_Ear2_Indel_R1 (WT yields 1339 bp amplicon) for session 1 with the 0:100 dCas9:Cas9 ratio. All amplicons were sent for Sanger sequencing. L1 = 1 kb DNA molecular weight ladder (thick band is 3 kb) and L2 = 100 bp DNA molecular weight ladder (thick band is 1kb). (c) Genotype outcomes for *Ear1* and *Ear2* for pups analysed and (d) summary table of analysis for electroporation and subsequent G0 animals. The alleles seen in each pup were put into the groupings of: unmodified animals indicating no mutation event for *Ear1*, partial wild type where the wild-type allele was seen in the presence of a mutated allele for *Ear1*, incorrect allele where a mutated allele was present that was not desired for *Ear1* and correct allele where a mutated allele was present and a desired allele for *Ear1*. * denotes animals with evidence of allele type.

Supplementary Figure 13: Analysis of the electroporation sessions for the *Ear2* project.

(a)The figure shows the PCR amplification of the *Ear1* paralog genomic region of interest with Geno_Ear1_Indel_F1 and Geno_Ear1_Indel_R1 (WT yields 1842 bp amplicon) for session 1 with the 0:100 dCas9:Cas9 ratio and (b) the PCR amplification of the *Ear2* genomic region of interest with Geno_Ear2_Indel_F1 and Geno_Ear2_Indel_R1 (WT yields 1339 bp amplicon) for session 1 with the 0:100 dCas9:Cas9 ratio. All amplicons were sent for Sanger sequencing. L1 = 1 kb DNA molecular weight ladder (thick band is 3 kb) and L2 = 100 bp DNA molecular weight ladder (thick band is 1kb). (c) Genotype outcomes for *Ear1* and *Ear2* for pups analysed and (d) summary table of analysis for electroporation and subsequent G0 animals. The alleles seen in each pup were put into the groupings of: unmodified animals indicating no mutation event for *Ear2*, partial wild type where the wild-type allele was seen in the presence of a mutated allele for *Ear2*, incorrect allele where a mutated allele was present that was not desired for *Ear2* and correct allele where a mutated allele was present and a desired allele for *Ear2*. * denotes animals with evidence of allele type.

