## Supplementary Figures for "Combining Cas9 and dCas9 facilitates genome editing in genes associated with viability or welfare issues, or within paralogous gene clusters"

### Slide 1
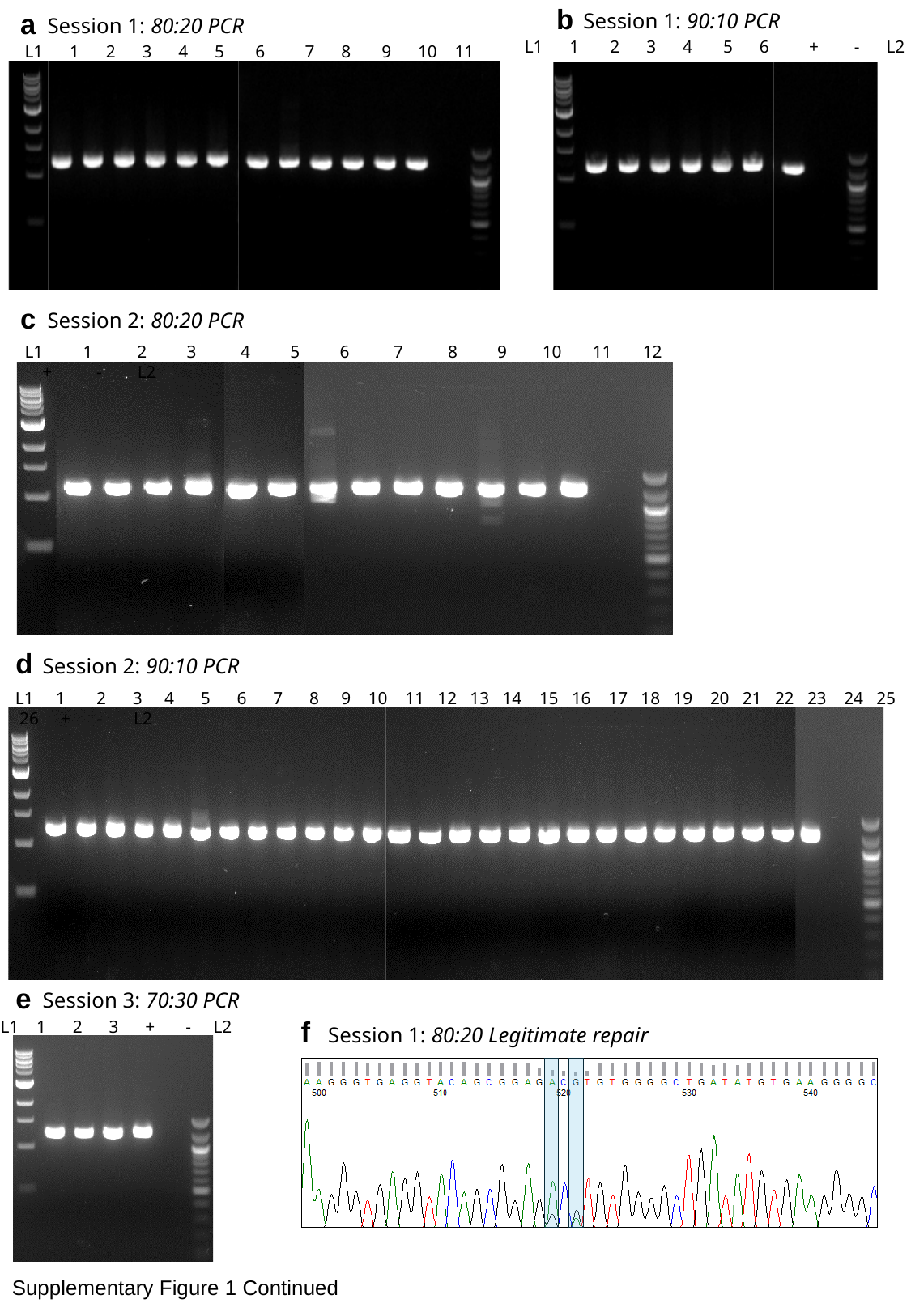

a
Session 1: 80:20 PCR
 L1 1 2 3 4 5 6 7 8 9 10 11 + - L2
b
Session 1: 90:10 PCR
L1 1 2 3 4 5 6 + - L2
c
Session 2: 80:20 PCR
 L1 1 2 3 4 5 6 7 8 9 10 11 12 + - L2
d
Session 2: 90:10 PCR
 L1 1 2 3 4 5 6 7 8 9 10 11 12 13 14 15 16 17 18 19 20 21 22 23 24 25 26 + - L2
e
Session 3: 70:30 PCR
L1 1 2 3 + - L2
f
Session 1: 80:20 Legitimate repair
Supplementary Figure 1 Continued

### Slide 2
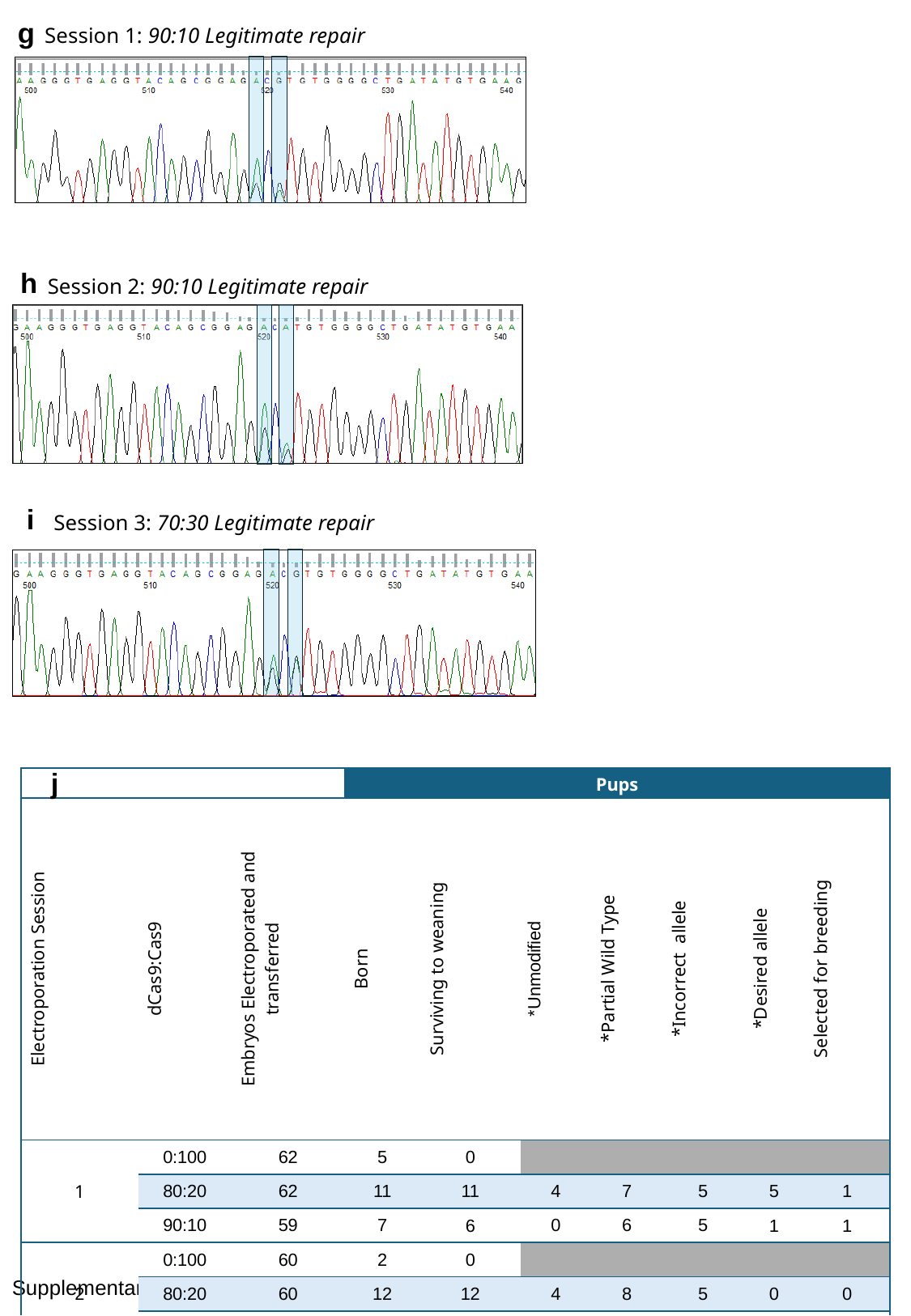

g
Session 1: 90:10 Legitimate repair
h
Session 2: 90:10 Legitimate repair
i
Session 3: 70:30 Legitimate repair
j
| | | | Pups | | | | | | |
| --- | --- | --- | --- | --- | --- | --- | --- | --- | --- |
| Electroporation Session | dCas9:Cas9 | Embryos Electroporated and transferred | Born | Surviving to weaning | \*Unmodified | \*Partial Wild Type | \*Incorrect allele | \*Desired allele | Selected for breeding |
| 1 | 0:100 | 62 | 5 | 0 | | | | | |
| | 80:20 | 62 | 11 | 11 | 4 | 7 | 5 | 5 | 1 |
| | 90:10 | 59 | 7 | 6 | 0 | 6 | 5 | 1 | 1 |
| 2 | 0:100 | 60 | 2 | 0 | | | | | |
| | 80:20 | 60 | 12 | 12 | 4 | 8 | 5 | 0 | 0 |
| | 90:10 | 60 | 27 | 26 | 10 | 16 | 15 | 7 | 0 |
| 3 | 0:100 | 80 | 0 | | | | | | |
| | 70:30 | 80 | 5 | 3 | 0 | 3 | 3 | 3 | 0 |
| | 80:20 | 80 | 0 | | | | | | |
Supplementary Figure 1

### Slide 3
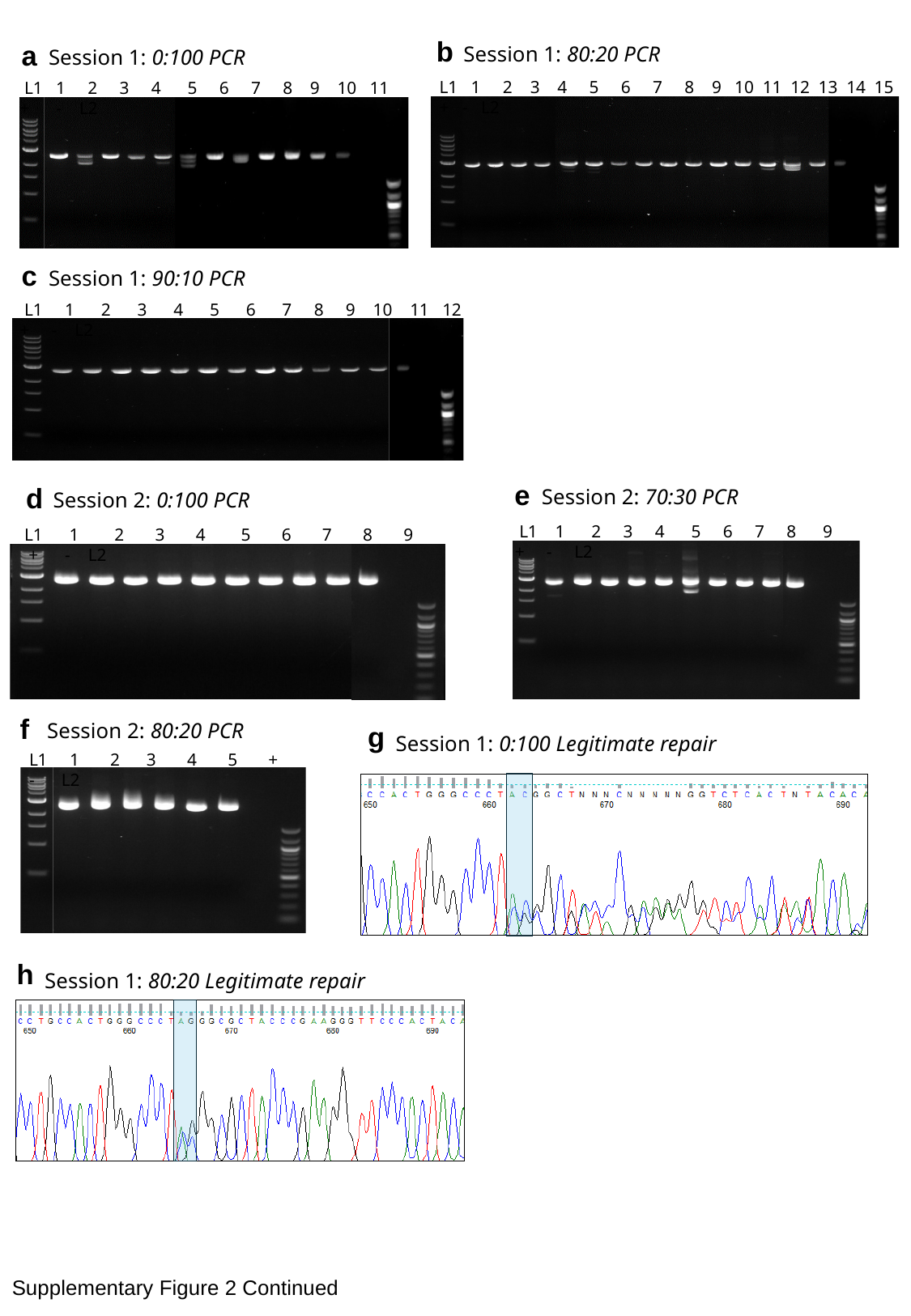

b
Session 1: 80:20 PCR
 L1 1 2 3 4 5 6 7 8 9 10 11 12 13 14 15 + - L2
a
Session 1: 0:100 PCR
 L1 1 2 3 4 5 6 7 8 9 10 11 + - L2
c
Session 1: 90:10 PCR
 L1 1 2 3 4 5 6 7 8 9 10 11 12 + - L2
e
Session 2: 70:30 PCR
 L1 1 2 3 4 5 6 7 8 9 + - L2
d
Session 2: 0:100 PCR
 L1 1 2 3 4 5 6 7 8 9 + - L2
f
Session 2: 80:20 PCR
 L1 1 2 3 4 5 + - L2
g
Session 1: 0:100 Legitimate repair
h
Session 1: 80:20 Legitimate repair
Supplementary Figure 2 Continued

### Slide 4
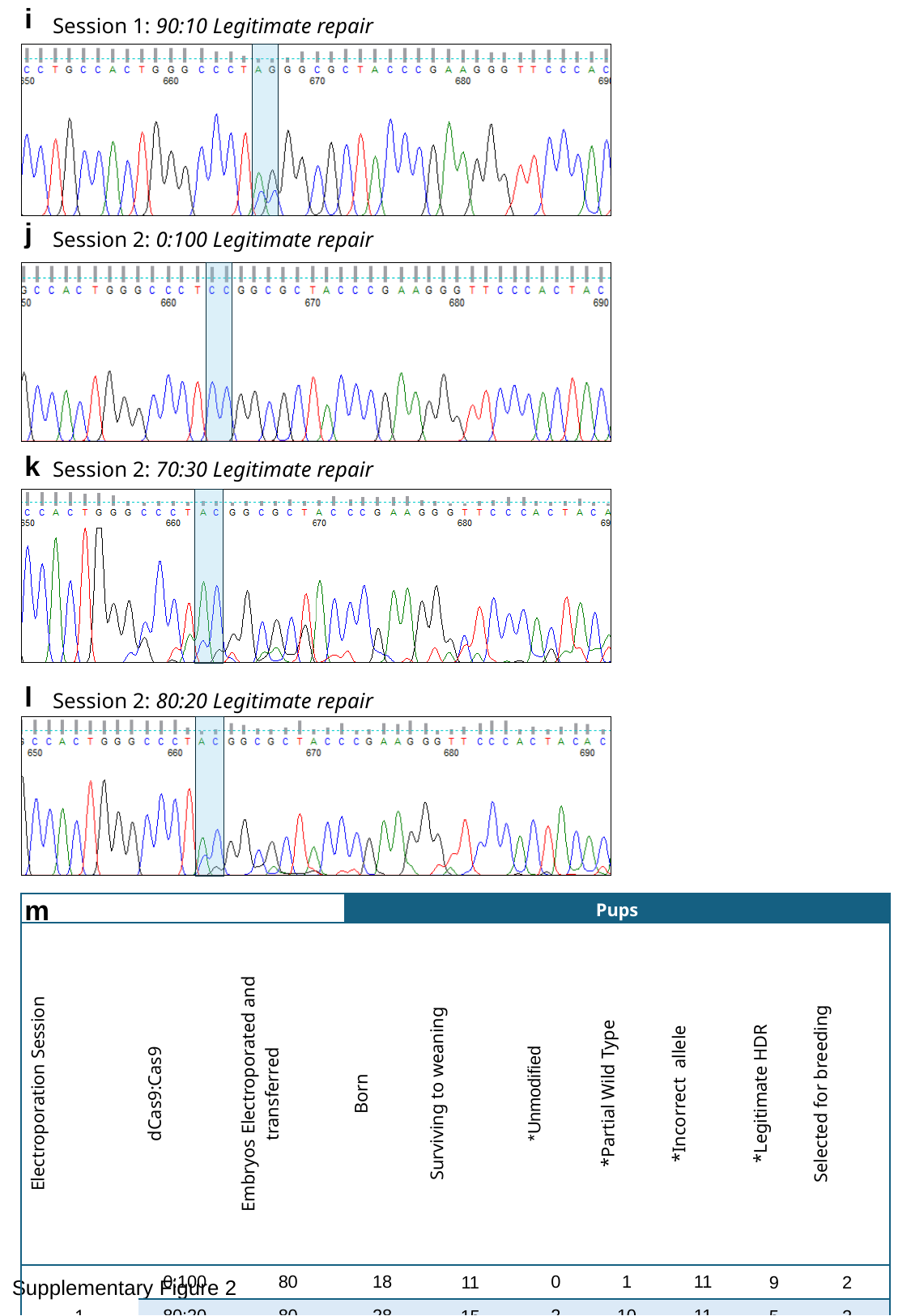

i
Session 1: 90:10 Legitimate repair
j
Session 2: 0:100 Legitimate repair
k
Session 2: 70:30 Legitimate repair
l
Session 2: 80:20 Legitimate repair
m
| | | | Pups | | | | | | |
| --- | --- | --- | --- | --- | --- | --- | --- | --- | --- |
| Electroporation Session | dCas9:Cas9 | Embryos Electroporated and transferred | Born | Surviving to weaning | \*Unmodified | \*Partial Wild Type | \*Incorrect allele | \*Legitimate HDR | Selected for breeding |
| 1 | 0:100 | 80 | 18 | 11 | 0 | 1 | 11 | 9 | 2 |
| | 80:20 | 80 | 28 | 15 | 2 | 10 | 11 | 5 | 2 |
| | 90:10 | 80 | 16 | 12 | 9 | 2 | 2 | 2 | 0 |
| 2 | 0:100 | 80 | 11 | 9 | 0 | 0 | 6 | 9 | 0 |
| | 70:30 | 80 | 10 | 9 | 3 | 5 | 6 | 2 | 0 |
| | 80:20 | 80 | 10 | 6^†^ | 1 | 3 | 4 | 2 | 0 |
Supplementary Figure 2

### Slide 5
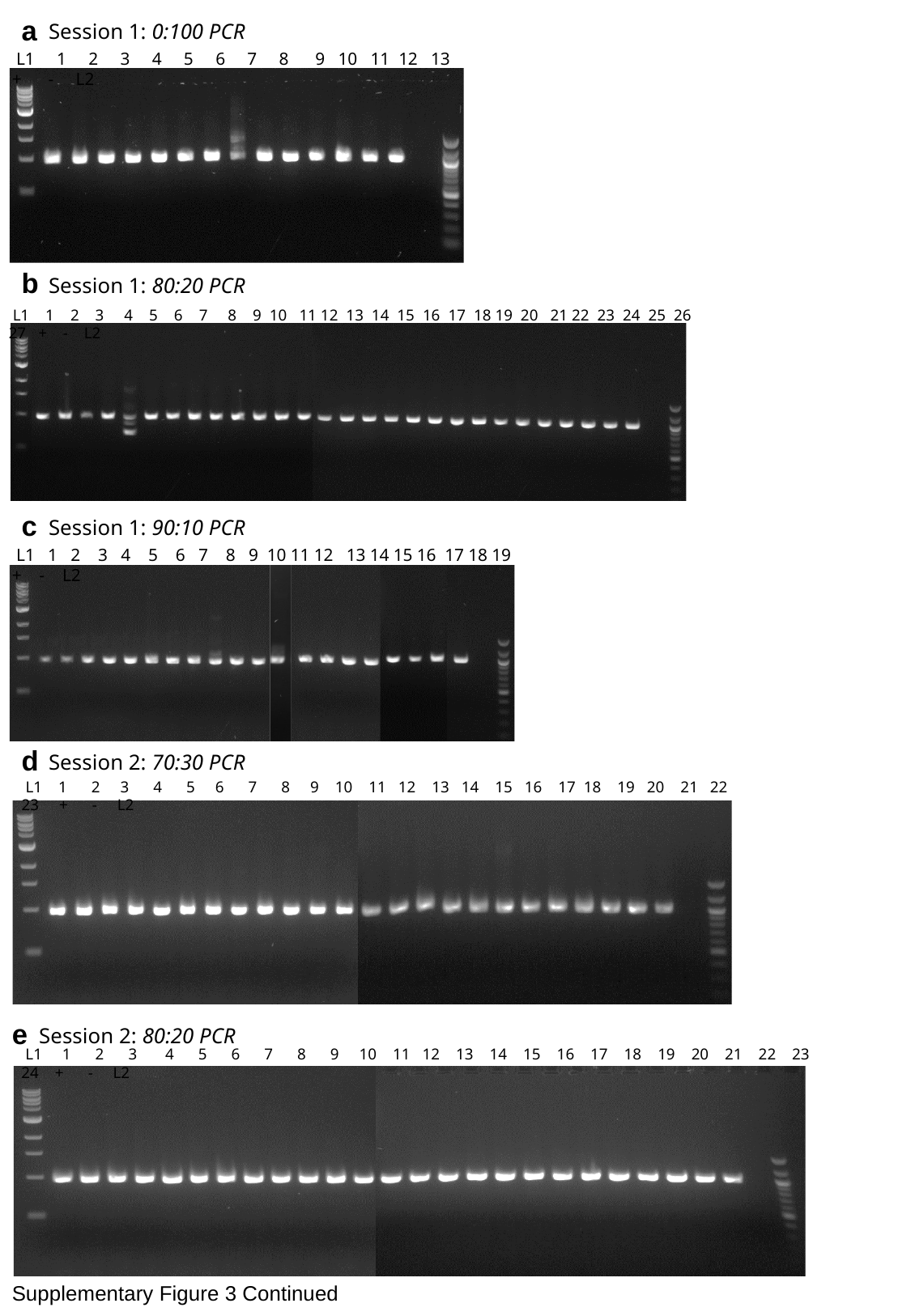

a
Session 1: 0:100 PCR
 L1 1 2 3 4 5 6 7 8 9 10 11 12 13 + - L2
b
Session 1: 80:20 PCR
 L1 1 2 3 4 5 6 7 8 9 10 11 12 13 14 15 16 17 18 19 20 21 22 23 24 25 26 27 + - L2
c
Session 1: 90:10 PCR
 L1 1 2 3 4 5 6 7 8 9 10 11 12 13 14 15 16 17 18 19 + - L2
d
Session 2: 70:30 PCR
 L1 1 2 3 4 5 6 7 8 9 10 11 12 13 14 15 16 17 18 19 20 21 22 23 + - L2
e
Session 2: 80:20 PCR
 L1 1 2 3 4 5 6 7 8 9 10 11 12 13 14 15 16 17 18 19 20 21 22 23 24 + - L2
Supplementary Figure 3 Continued

### Slide 6
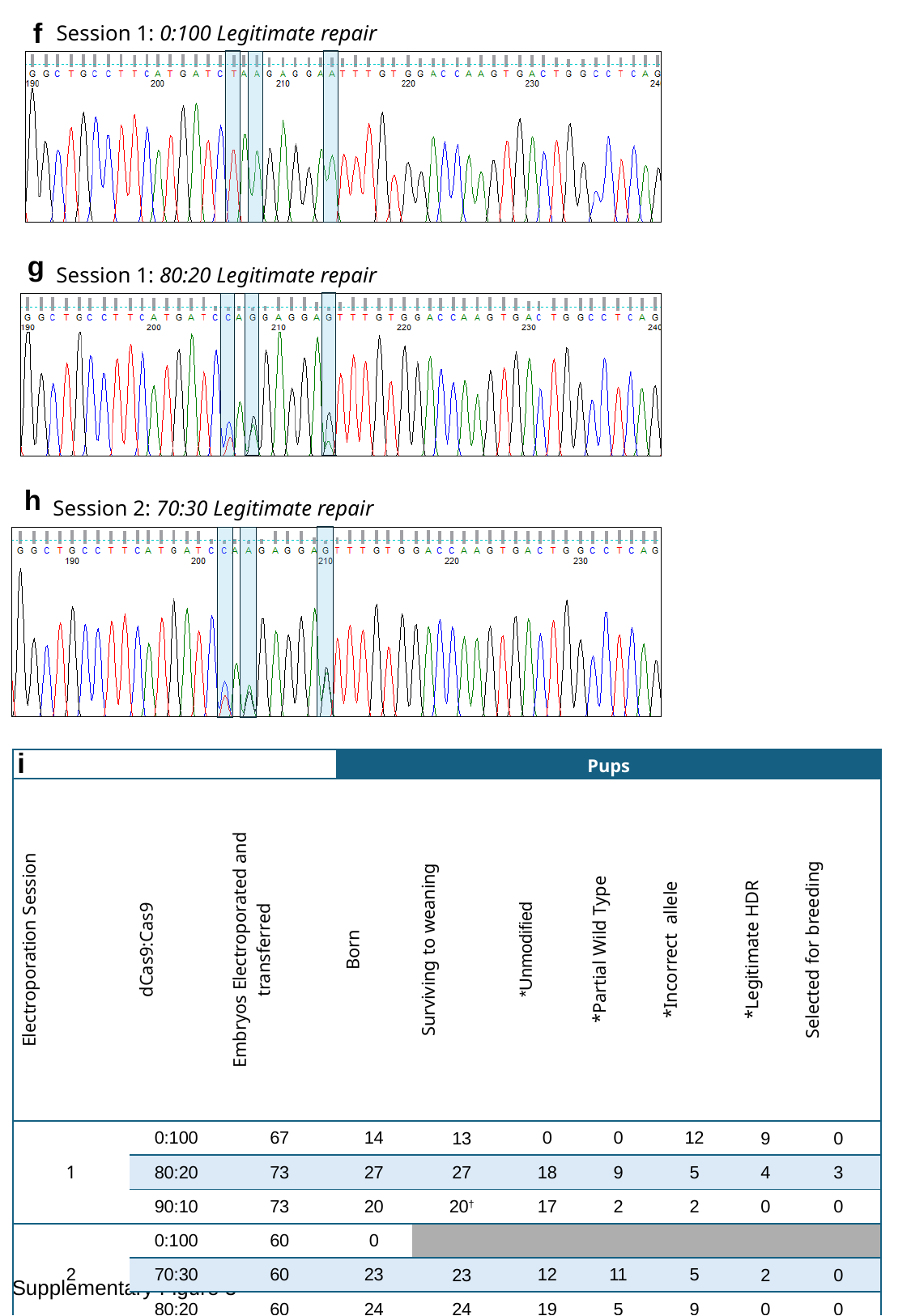

f
Session 1: 0:100 Legitimate repair
g
Session 1: 80:20 Legitimate repair
h
Session 2: 70:30 Legitimate repair
i
| | | | Pups | | | | | | |
| --- | --- | --- | --- | --- | --- | --- | --- | --- | --- |
| Electroporation Session | dCas9:Cas9 | Embryos Electroporated and transferred | Born | Surviving to weaning | \*Unmodified | \*Partial Wild Type | \*Incorrect allele | \*Legitimate HDR | Selected for breeding |
| 1 | 0:100 | 67 | 14 | 13 | 0 | 0 | 12 | 9 | 0 |
| | 80:20 | 73 | 27 | 27 | 18 | 9 | 5 | 4 | 3 |
| | 90:10 | 73 | 20 | 20^†^ | 17 | 2 | 2 | 0 | 0 |
| 2 | 0:100 | 60 | 0 | | | | | | |
| | 70:30 | 60 | 23 | 23 | 12 | 11 | 5 | 2 | 0 |
| | 80:20 | 60 | 24 | 24 | 19 | 5 | 9 | 0 | 0 |
Supplementary Figure 3

### Slide 7
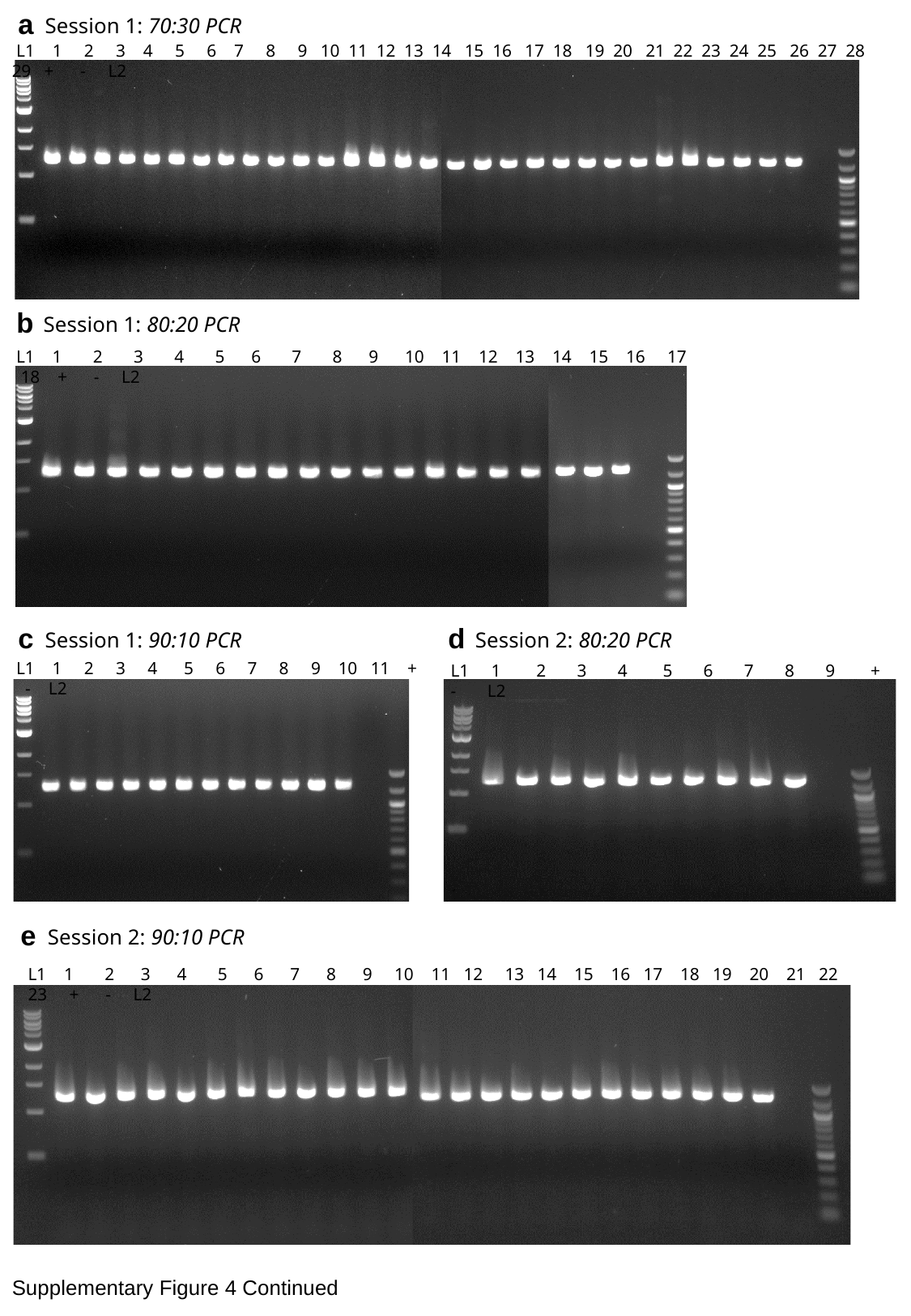

a
Session 1: 70:30 PCR
 L1 1 2 3 4 5 6 7 8 9 10 11 12 13 14 15 16 17 18 19 20 21 22 23 24 25 26 27 28 29 + - L2
b
Session 1: 80:20 PCR
 L1 1 2 3 4 5 6 7 8 9 10 11 12 13 14 15 16 17 18 + - L2
c
Session 1: 90:10 PCR
 L1 1 2 3 4 5 6 7 8 9 10 11 + - L2
d
Session 2: 80:20 PCR
L1 1 2 3 4 5 6 7 8 9 + - L2
e
Session 2: 90:10 PCR
 L1 1 2 3 4 5 6 7 8 9 10 11 12 13 14 15 16 17 18 19 20 21 22 23 + - L2
Supplementary Figure 4 Continued

### Slide 8
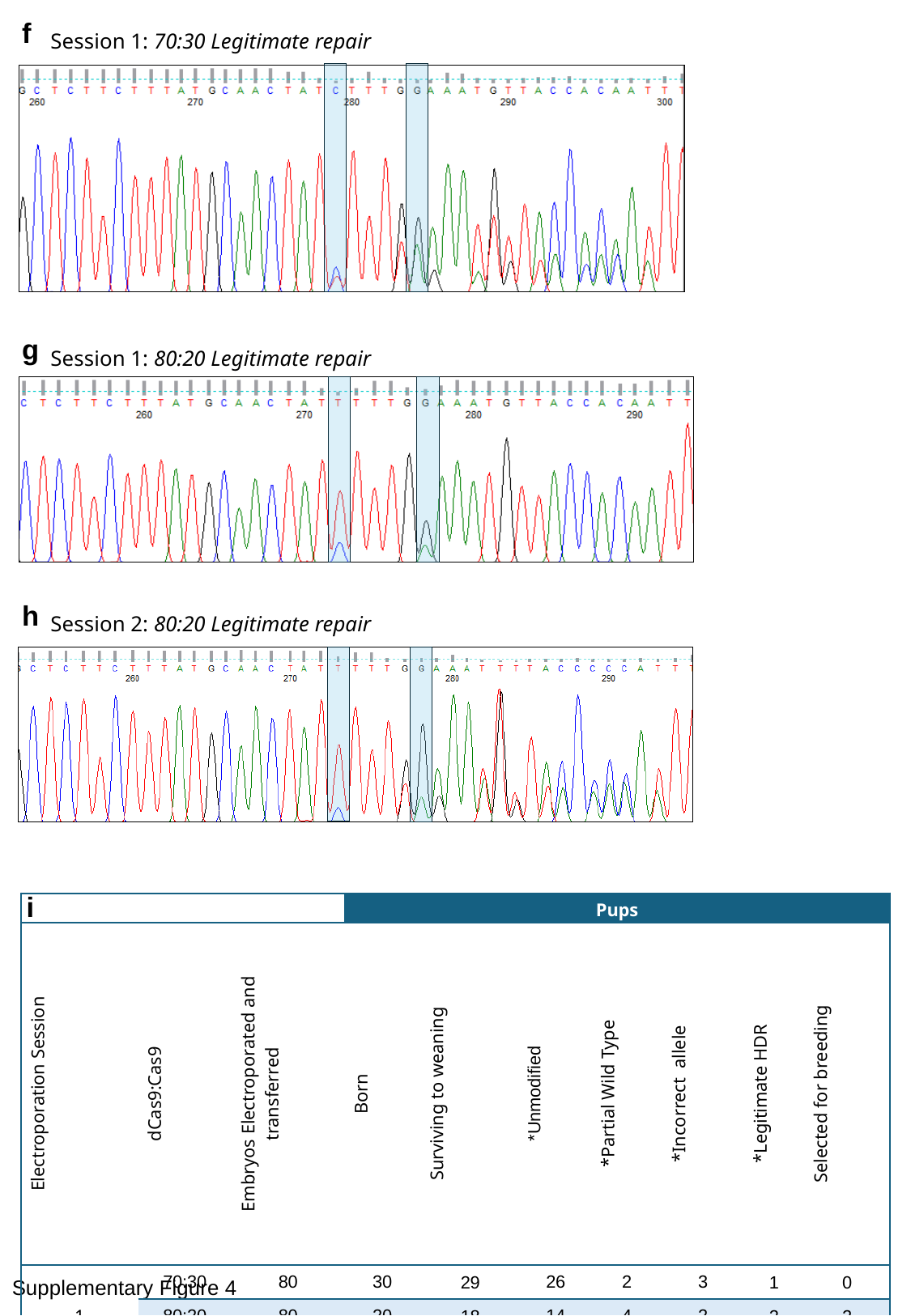

f
Session 1: 70:30 Legitimate repair
g
Session 1: 80:20 Legitimate repair
h
Session 2: 80:20 Legitimate repair
i
| | | | Pups | | | | | | |
| --- | --- | --- | --- | --- | --- | --- | --- | --- | --- |
| Electroporation Session | dCas9:Cas9 | Embryos Electroporated and transferred | Born | Surviving to weaning | \*Unmodified | \*Partial Wild Type | \*Incorrect allele | \*Legitimate HDR | Selected for breeding |
| 1 | 70:30 | 80 | 30 | 29 | 26 | 2 | 3 | 1 | 0 |
| | 80:20 | 80 | 20 | 18 | 14 | 4 | 2 | 2 | 2 |
| | 90:10 | 80 | 18 | 11 | 9 | 2 | 2 | 0 | 0 |
| 2 | 80:20 | 80 | 11 | 10^†^ | 4 | 4 | 5 | 1 | 1 |
| | 90:10 | 80 | 24 | 23 | 15 | 8 | 8 | 0 | 0 |
Supplementary Figure 4

### Slide 9
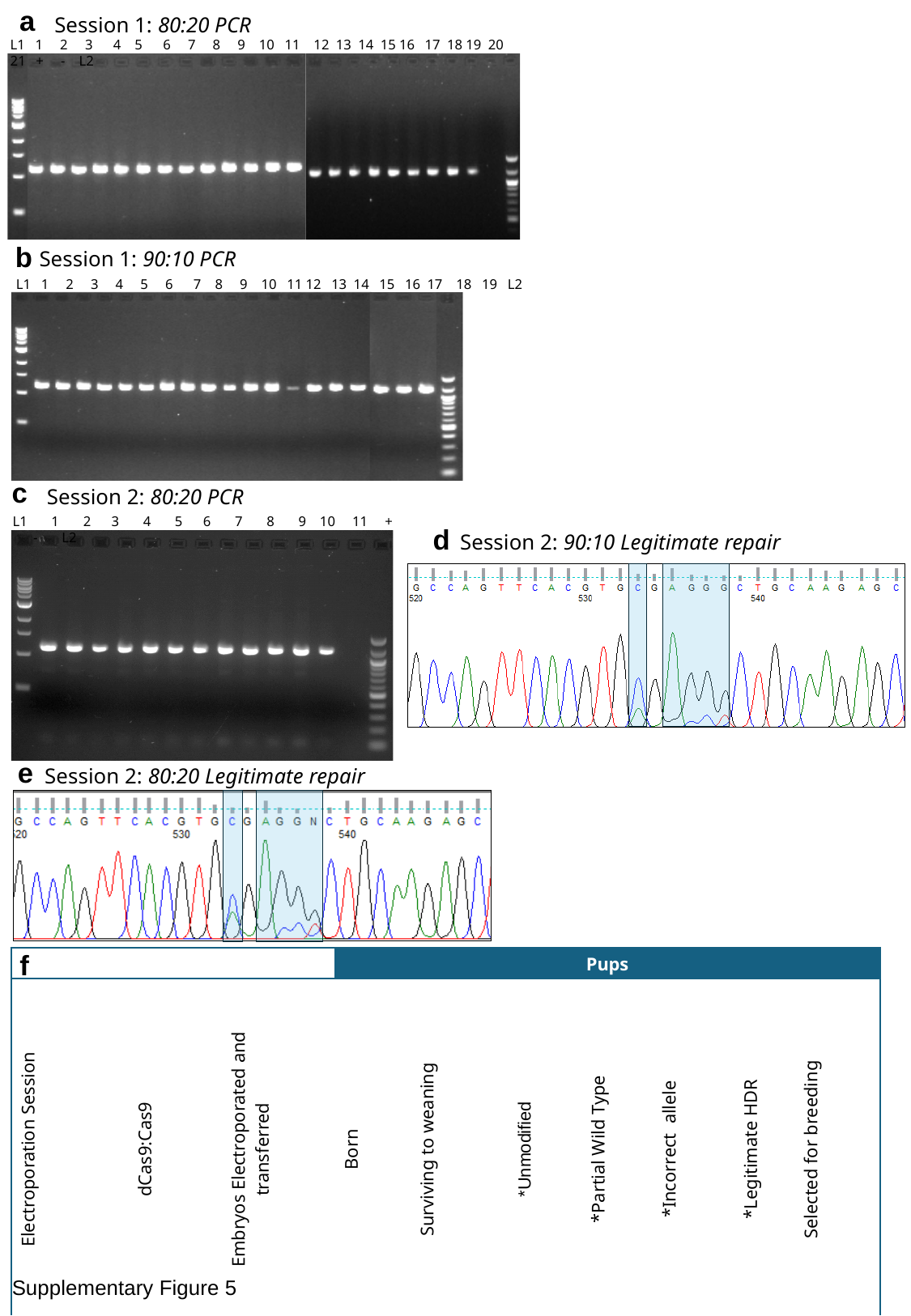

a
Session 1: 80:20 PCR
L1 1 2 3 4 5 6 7 8 9 10 11 12 13 14 15 16 17 18 19 20 21 + - L2
b
Session 1: 90:10 PCR
L1 1 2 3 4 5 6 7 8 9 10 11 12 13 14 15 16 17 18 19 L2
c
Session 2: 80:20 PCR
L1 1 2 3 4 5 6 7 8 9 10 11 + - L2
d
Session 2: 90:10 Legitimate repair
e
Session 2: 80:20 Legitimate repair
f
| | | | Pups | | | | | | |
| --- | --- | --- | --- | --- | --- | --- | --- | --- | --- |
| Electroporation Session | dCas9:Cas9 | Embryos Electroporated and transferred | Born | Surviving to weaning | \*Unmodified | \*Partial Wild Type | \*Incorrect allele | \*Legitimate HDR | Selected for breeding |
| 1 | 0:100 | 80 | 0 | | | | | | |
| | 80:20 | 80 | 21 | 21 | 17 | 4 | 4 | 0 | 0 |
| | 90:10 | 80 | 34 | 19 | 15 | 4 | 3 | 1 | 1 |
| 2 | 70:30 | 8 | 0 | | | | | | |
| | 80:20 | 80 | 12 | 11 | 5 | 6 | 5 | 1 | 1 |
Supplementary Figure 5

### Slide 10
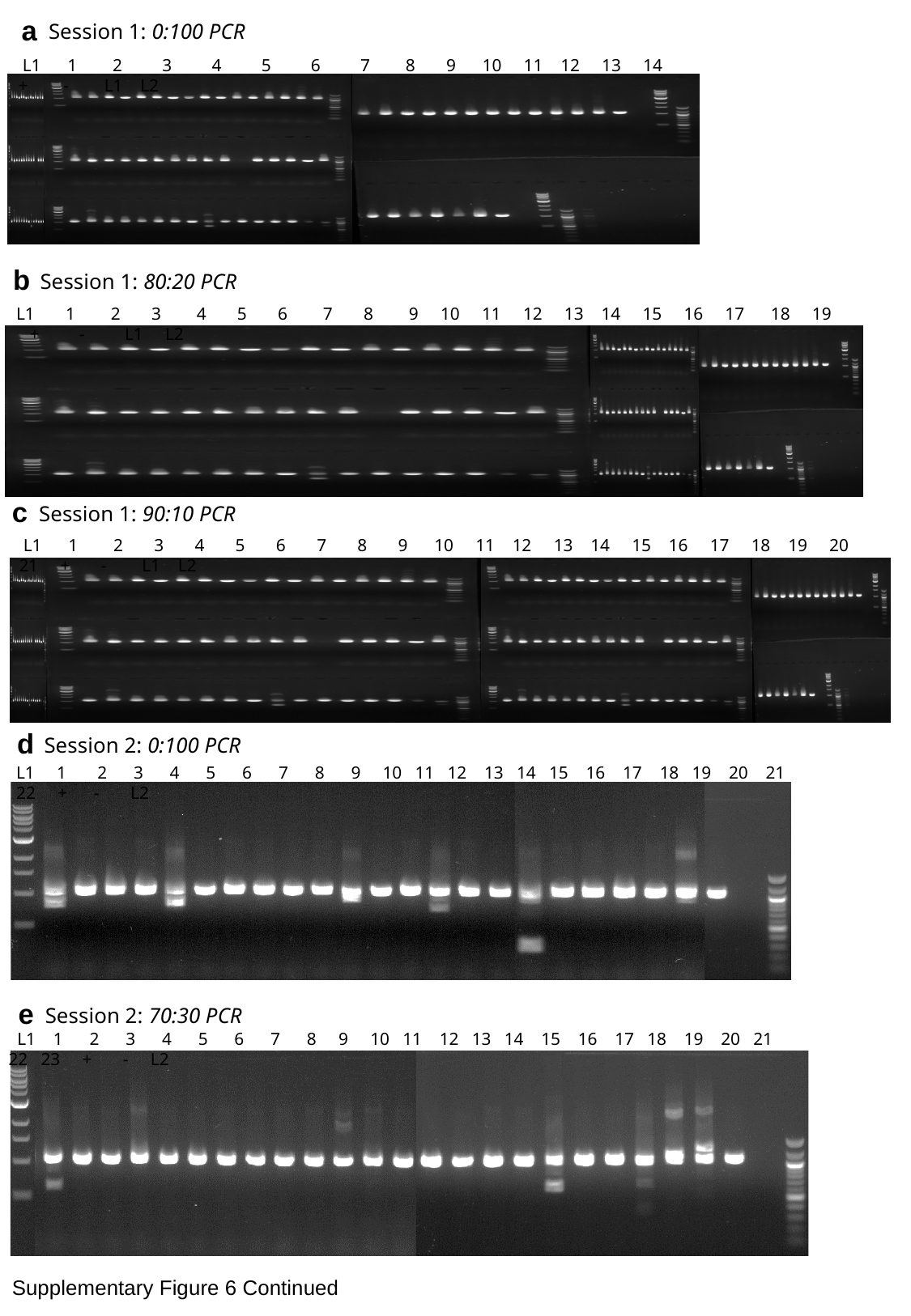

a
Session 1: 0:100 PCR
 L1 1 2 3 4 5 6 7 8 9 10 11 12 13 14 + - L1 L2
b
Session 1: 80:20 PCR
 L1 1 2 3 4 5 6 7 8 9 10 11 12 13 14 15 16 17 18 19 + - L1 L2
c
Session 1: 90:10 PCR
 L1 1 2 3 4 5 6 7 8 9 10 11 12 13 14 15 16 17 18 19 20 21 + - L1 L2
d
Session 2: 0:100 PCR
 L1 1 2 3 4 5 6 7 8 9 10 11 12 13 14 15 16 17 18 19 20 21 22 + - L2
e
Session 2: 70:30 PCR
 L1 1 2 3 4 5 6 7 8 9 10 11 12 13 14 15 16 17 18 19 20 21 22 23 + - L2
Supplementary Figure 6 Continued

### Slide 11
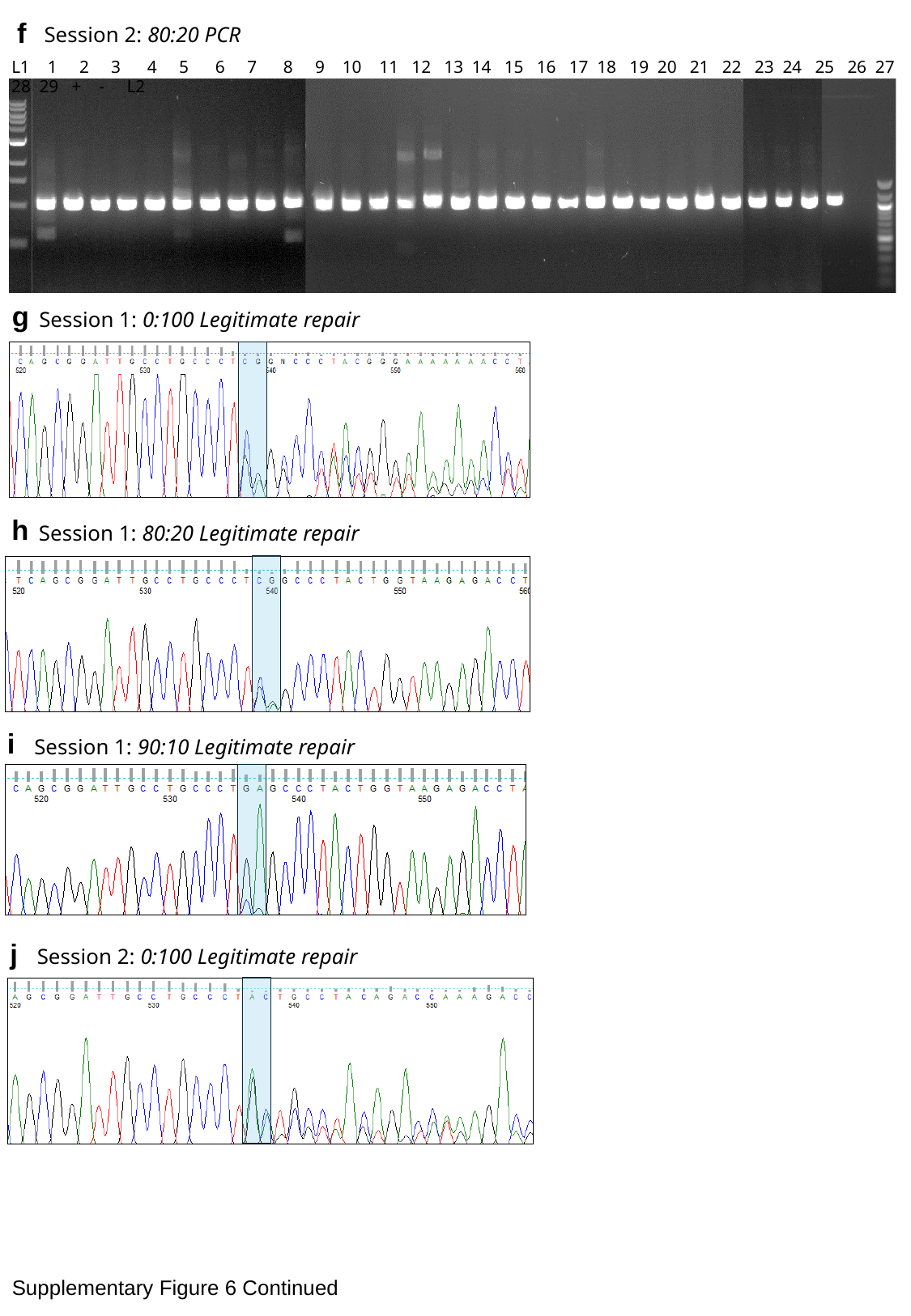

f
Session 2: 80:20 PCR
 L1 1 2 3 4 5 6 7 8 9 10 11 12 13 14 15 16 17 18 19 20 21 22 23 24 25 26 27 28 29 + - L2
g
Session 1: 0:100 Legitimate repair
h
Session 1: 80:20 Legitimate repair
i
Session 1: 90:10 Legitimate repair
j
Session 2: 0:100 Legitimate repair
Supplementary Figure 6 Continued

### Slide 12
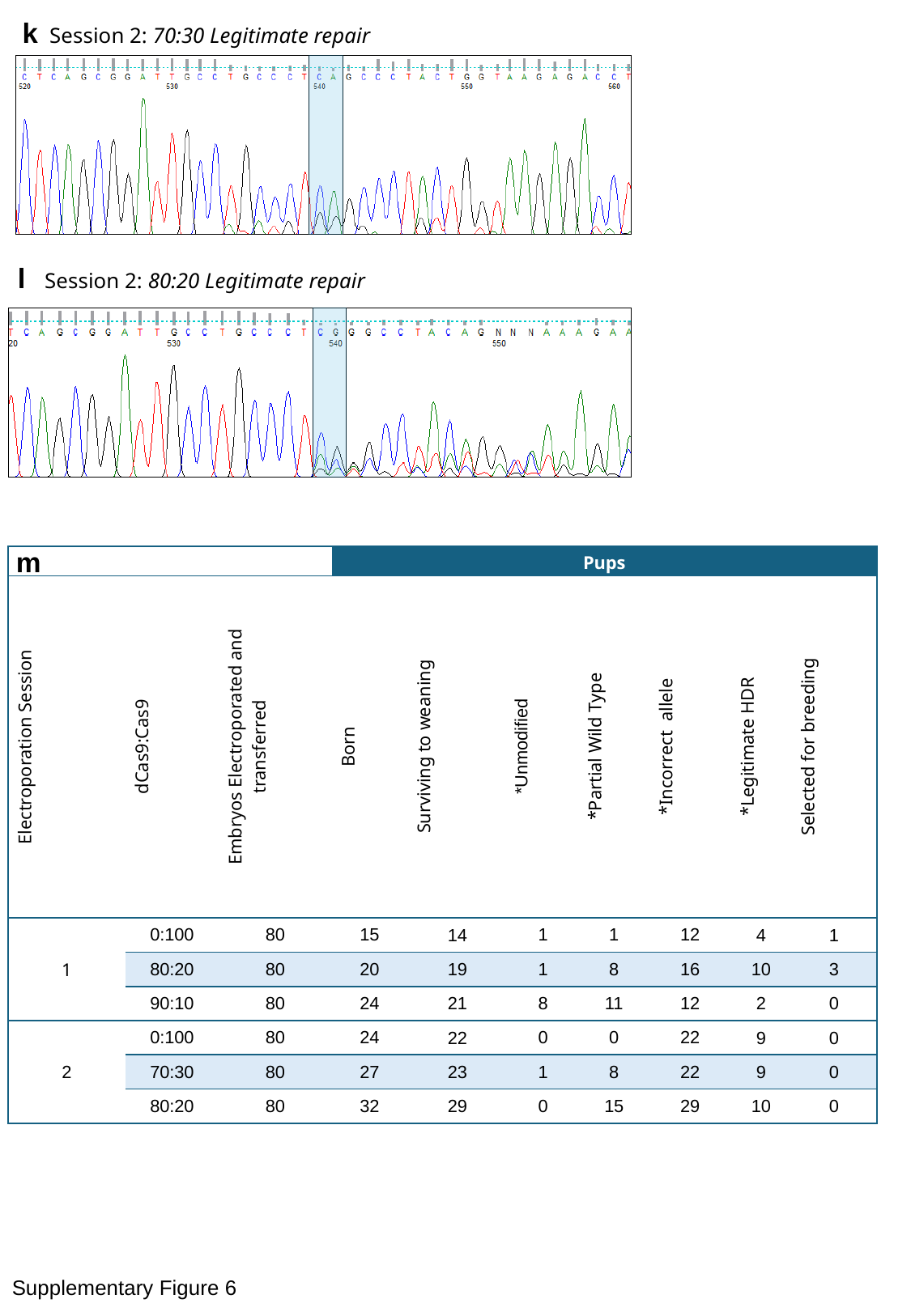

k
Session 2: 70:30 Legitimate repair
l
Session 2: 80:20 Legitimate repair
m
| | | | Pups | | | | | | |
| --- | --- | --- | --- | --- | --- | --- | --- | --- | --- |
| Electroporation Session | dCas9:Cas9 | Embryos Electroporated and transferred | Born | Surviving to weaning | \*Unmodified | \*Partial Wild Type | \*Incorrect allele | \*Legitimate HDR | Selected for breeding |
| 1 | 0:100 | 80 | 15 | 14 | 1 | 1 | 12 | 4 | 1 |
| | 80:20 | 80 | 20 | 19 | 1 | 8 | 16 | 10 | 3 |
| | 90:10 | 80 | 24 | 21 | 8 | 11 | 12 | 2 | 0 |
| 2 | 0:100 | 80 | 24 | 22 | 0 | 0 | 22 | 9 | 0 |
| | 70:30 | 80 | 27 | 23 | 1 | 8 | 22 | 9 | 0 |
| | 80:20 | 80 | 32 | 29 | 0 | 15 | 29 | 10 | 0 |
Supplementary Figure 6

### Slide 13
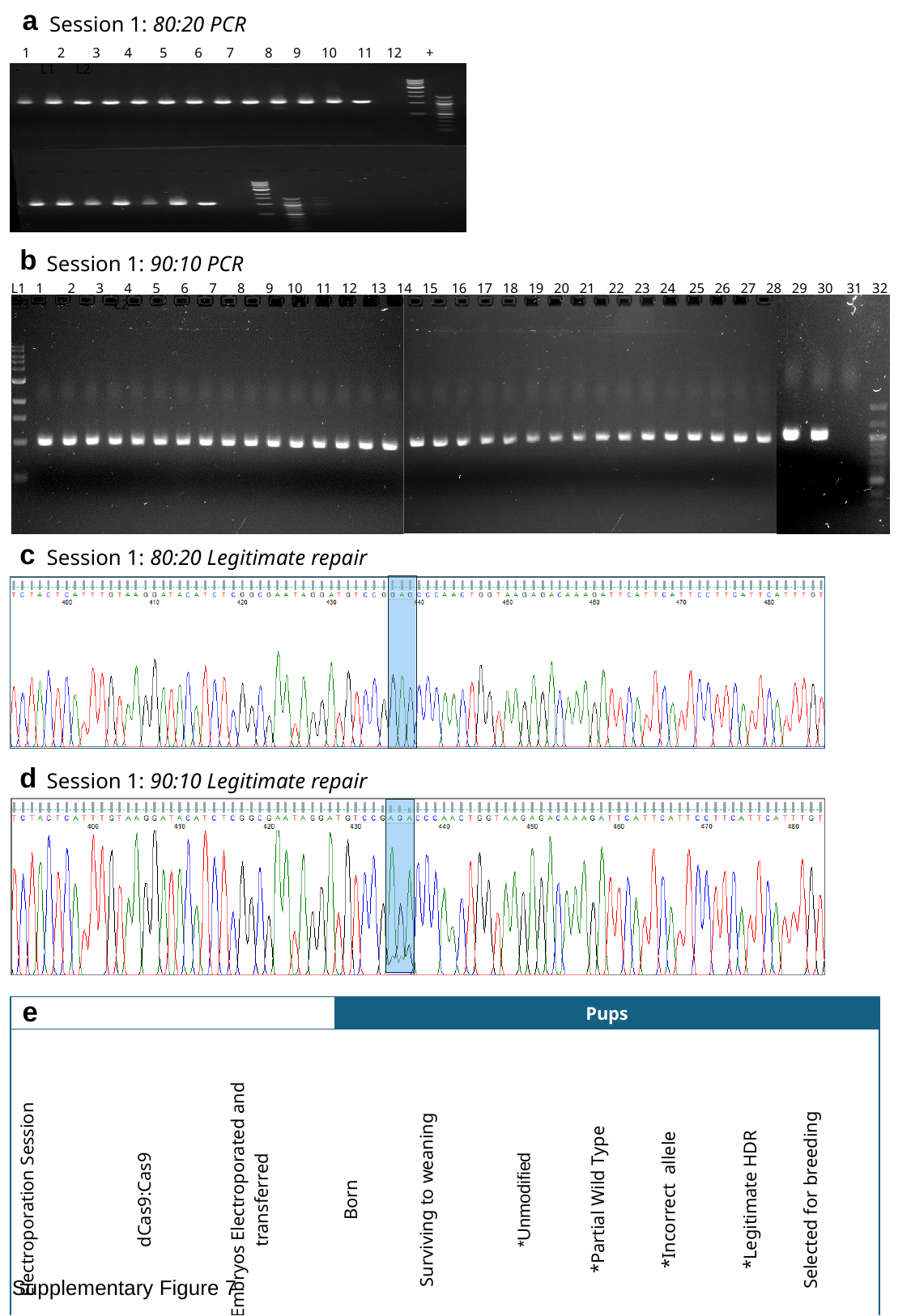

a
Session 1: 80:20 PCR
 1 2 3 4 5 6 7 8 9 10 11 12 + - L1 L2
b
Session 1: 90:10 PCR
L1 1 2 3 4 5 6 7 8 9 10 11 12 13 14 15 16 17 18 19 20 21 22 23 24 25 26 27 28 29 30 31 32 33 + - L2
c
Session 1: 80:20 Legitimate repair
d
Session 1: 90:10 Legitimate repair
e
| | | | Pups | | | | | | |
| --- | --- | --- | --- | --- | --- | --- | --- | --- | --- |
| Electroporation Session | dCas9:Cas9 | Embryos Electroporated and transferred | Born | Surviving to weaning | \*Unmodified | \*Partial Wild Type | \*Incorrect allele | \*Legitimate HDR | Selected for breeding |
| 1 | 0:100 | 80 | 4 | 0 | | | | | |
| | 80:20 | 80 | 16 | 12 | 3 | 5 | 8 | 1 | 1 |
| | 90:10 | 80 | 34 | 33 | 21 | 12 | 10 | 2 | 2 |
Supplementary Figure 7

### Slide 14
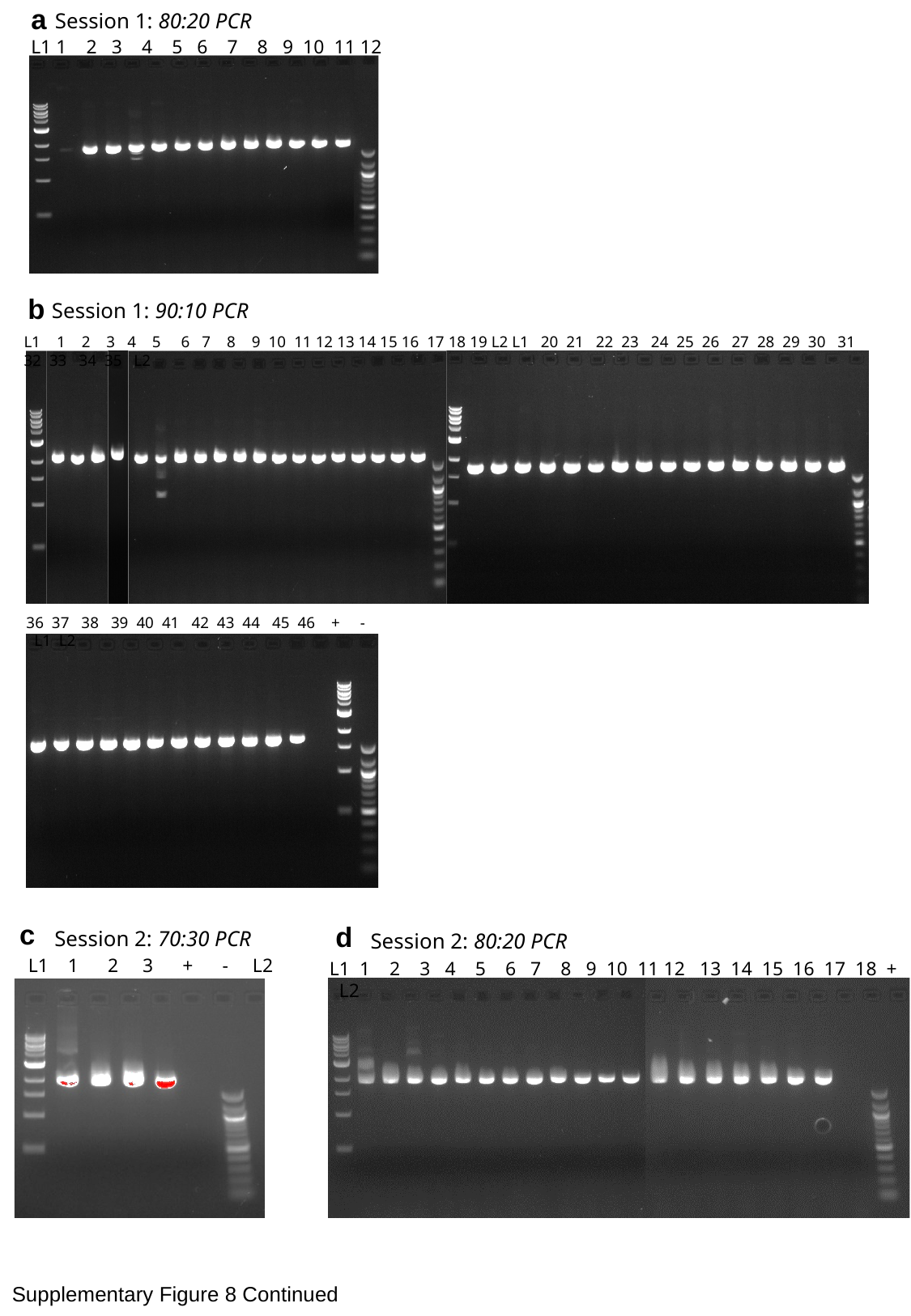

a
Session 1: 80:20 PCR
L1 1 2 3 4 5 6 7 8 9 10 11 12 13 L2
b
Session 1: 90:10 PCR
L1 1 2 3 4 5 6 7 8 9 10 11 12 13 14 15 16 17 18 19 L2 L1 20 21 22 23 24 25 26 27 28 29 30 31 32 33 34 35 L2
36 37 38 39 40 41 42 43 44 45 46 + - L1 L2
c
Session 2: 70:30 PCR
 L1 1 2 3 + - L2
d
Session 2: 80:20 PCR
L1 1 2 3 4 5 6 7 8 9 10 11 12 13 14 15 16 17 18 + - L2
Supplementary Figure 8 Continued

### Slide 15
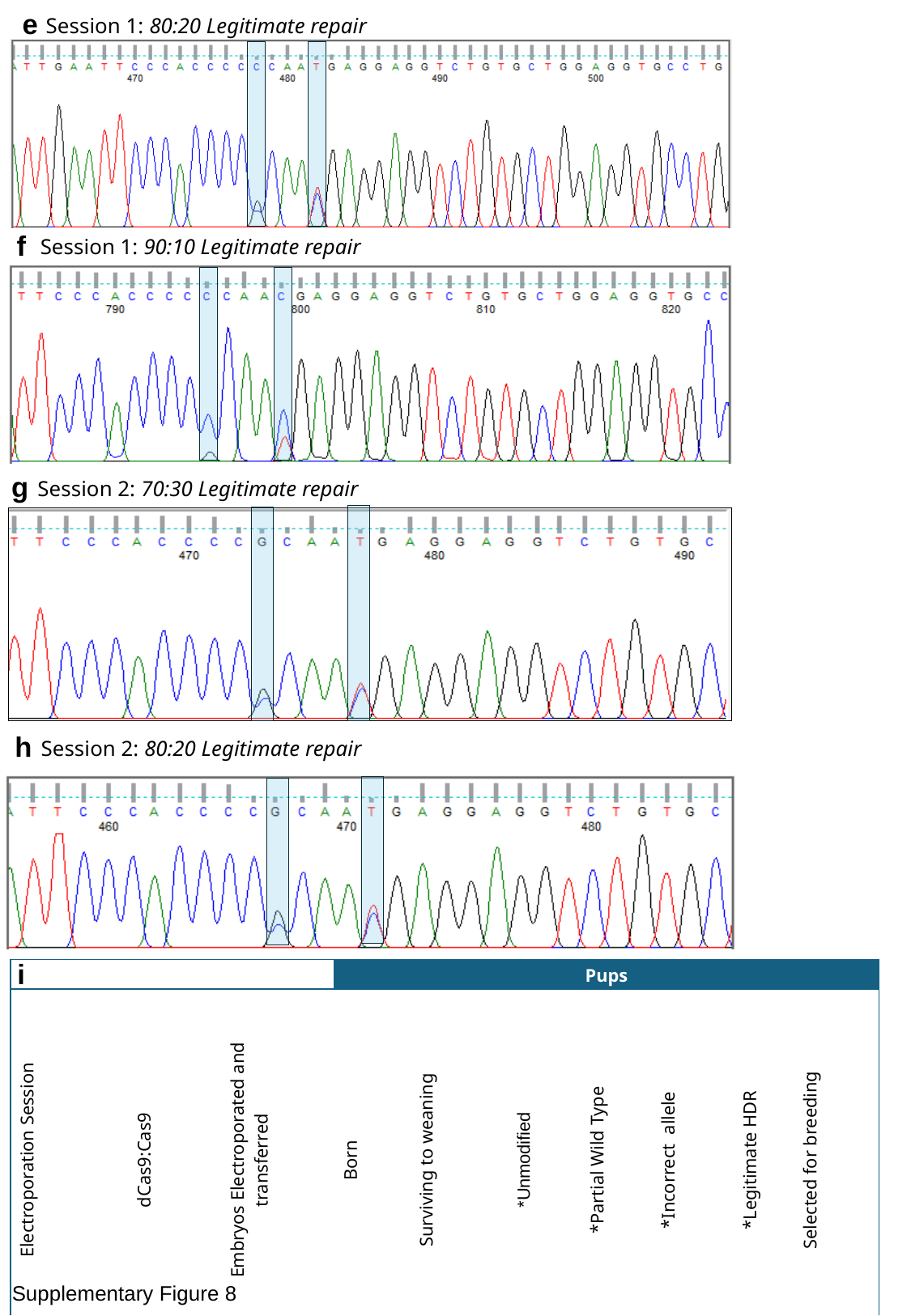

e
Session 1: 80:20 Legitimate repair
f
Session 1: 90:10 Legitimate repair
g
Session 2: 70:30 Legitimate repair
h
Session 2: 80:20 Legitimate repair
i
| | | | Pups | | | | | | |
| --- | --- | --- | --- | --- | --- | --- | --- | --- | --- |
| Electroporation Session | dCas9:Cas9 | Embryos Electroporated and transferred | Born | Surviving to weaning | \*Unmodified | \*Partial Wild Type | \*Incorrect allele | \*Legitimate HDR | Selected for breeding |
| 1 | 80:20 | 120 | 16 | 13 | 1 | 12 | 11 | 5 | 0 |
| | 90:10 | 120 | 50 | 46 | 27 | 19 | 12 | 7 | 4 |
| 2 | 0:100 | 80 | 0 | | | | | | |
| | 70:30 | 80 | 3 | 3 | 1 | 2 | 2 | 2 | 0 |
| | 80:20 | 80 | 19 | 19^†^ | 1 | 16 | 4 | 4 | 0 |
Supplementary Figure 8

### Slide 16
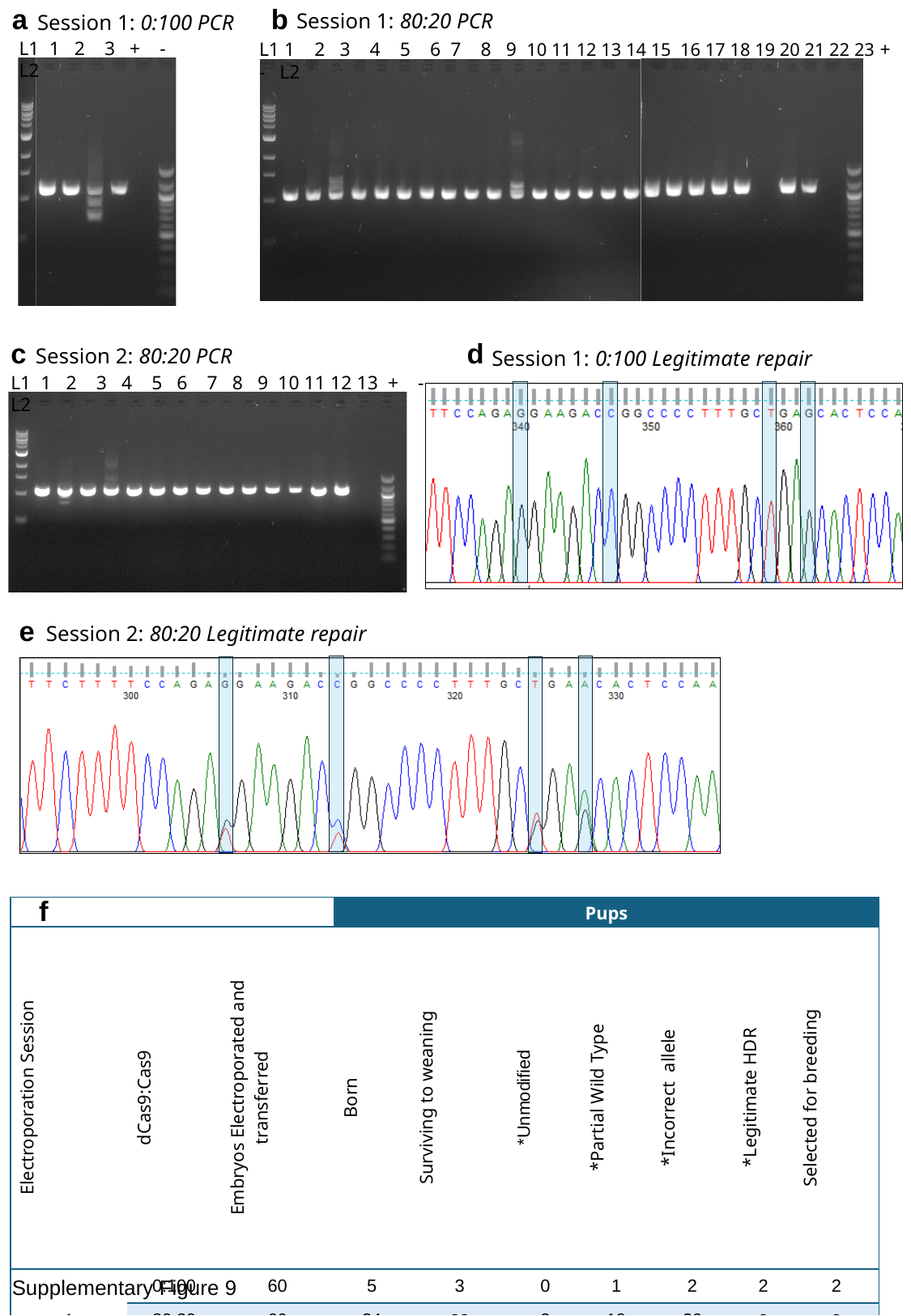

b
Session 1: 80:20 PCR
L1 1 2 3 4 5 6 7 8 9 10 11 12 13 14 15 16 17 18 19 20 21 22 23 + - L2
a
Session 1: 0:100 PCR
L1 1 2 3 + - L2
c
Session 2: 80:20 PCR
L1 1 2 3 4 5 6 7 8 9 10 11 12 13 + - L2
d
Session 1: 0:100 Legitimate repair
e
Session 2: 80:20 Legitimate repair
f
| | | | Pups | | | | | | |
| --- | --- | --- | --- | --- | --- | --- | --- | --- | --- |
| Electroporation Session | dCas9:Cas9 | Embryos Electroporated and transferred | Born | Surviving to weaning | \*Unmodified | \*Partial Wild Type | \*Incorrect allele | \*Legitimate HDR | Selected for breeding |
| 1 | 0:100 | 60 | 5 | 3 | 0 | 1 | 2 | 2 | 2 |
| | 80:20 | 60 | 24 | 23 | 2 | 19 | 20 | 2 | 2 |
| | 90:10 | 60 | 13 | 0 | | | | | |
| 2 | 0:100 | 80 | 0 | | | | | | |
| | 70:30 | 80 | 0 | | | | | | |
| | 80:20 | 80 | 13 | 13 | 2 | 5 | 10 | 3 | 0 |
Supplementary Figure 9

### Slide 17
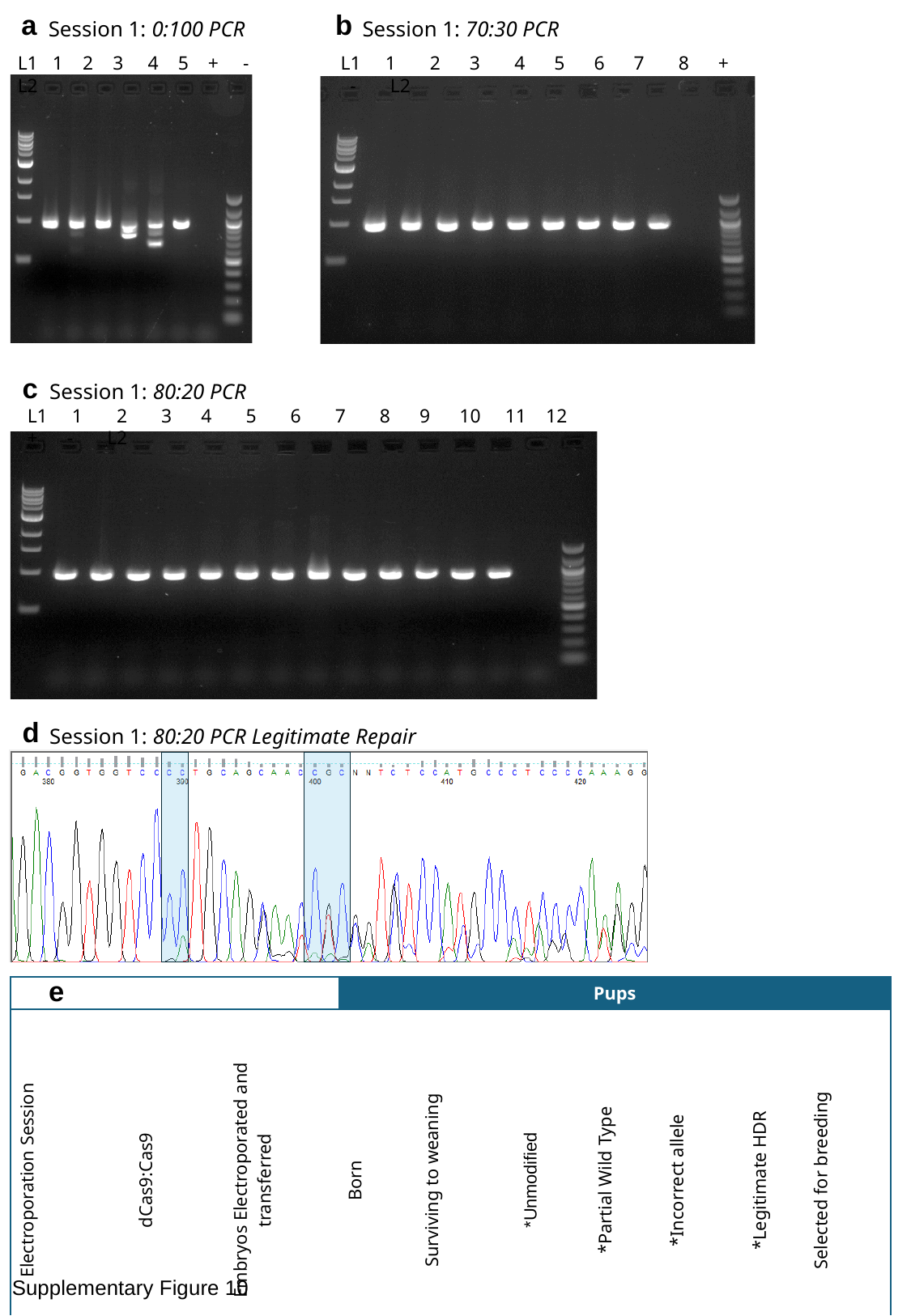

a
Session 1: 0:100 PCR
L1 1 2 3 4 5 + - L2
b
Session 1: 70:30 PCR
 L1 1 2 3 4 5 6 7 8 + - L2
c
Session 1: 80:20 PCR
 L1 1 2 3 4 5 6 7 8 9 10 11 12 + - L2
d
Session 1: 80:20 PCR Legitimate Repair
e
| | | | Pups | | | | | | |
| --- | --- | --- | --- | --- | --- | --- | --- | --- | --- |
| Electroporation Session | dCas9:Cas9 | Embryos Electroporated and transferred | Born | Surviving to weaning | \*Unmodified | \*Partial Wild Type | \*Incorrect allele | \*Legitimate HDR | Selected for breeding |
| 1 | 0:100 | 78 | 5 | 5 | 0 | 2 | 5 | 0 | 0 |
| | 70:30 | 80 | 15 | 8 | 6 | 2 | 2 | 0 | 0 |
| | 80:20 | 74 | 12 | 12 | 8 | 3 | 4 | 1 | 1 |
Supplementary Figure 10

### Slide 18
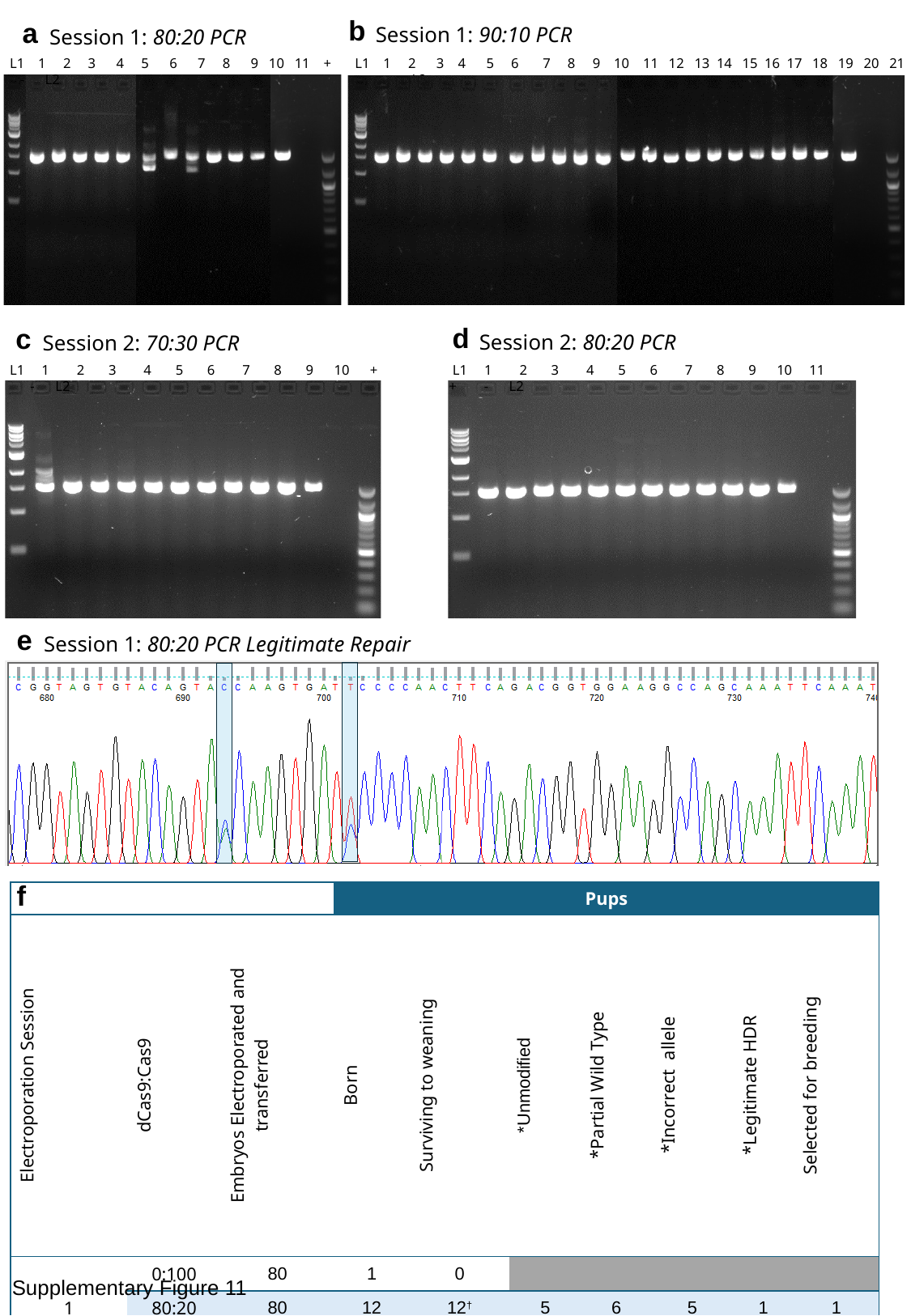

b
Session 1: 90:10 PCR
 L1 1 2 3 4 5 6 7 8 9 10 11 12 13 14 15 16 17 18 19 20 21 + - L2
a
Session 1: 80:20 PCR
 L1 1 2 3 4 5 6 7 8 9 10 11 + - L2
d
c
Session 2: 80:20 PCR
Session 2: 70:30 PCR
 L1 1 2 3 4 5 6 7 8 9 10 11 + - L2
 L1 1 2 3 4 5 6 7 8 9 10 + - L2
e
Session 1: 80:20 PCR Legitimate Repair
f
| | | | Pups | | | | | | |
| --- | --- | --- | --- | --- | --- | --- | --- | --- | --- |
| Electroporation Session | dCas9:Cas9 | Embryos Electroporated and transferred | Born | Surviving to weaning | \*Unmodified | \*Partial Wild Type | \*Incorrect allele | \*Legitimate HDR | Selected for breeding |
| 1 | 0:100 | 80 | 1 | 0 | | | | | |
| | 80:20 | 80 | 12 | 12^†^ | 5 | 6 | 5 | 1 | 1 |
| | 90:10 | 80 | 22 | 21 | 14 | 7 | 7 | 0 | 1^‡^ |
| 2 | 0:100 | 60 | 0 | | | | | | |
| | 70:30 | 60 | 10 | 10 | 1 | 8 | 9 | 0 | 0 |
| | 80:20 | 60 | 11 | 11 | 7 | 3 | 4 | 0 | 0 |
Supplementary Figure 11

### Slide 19
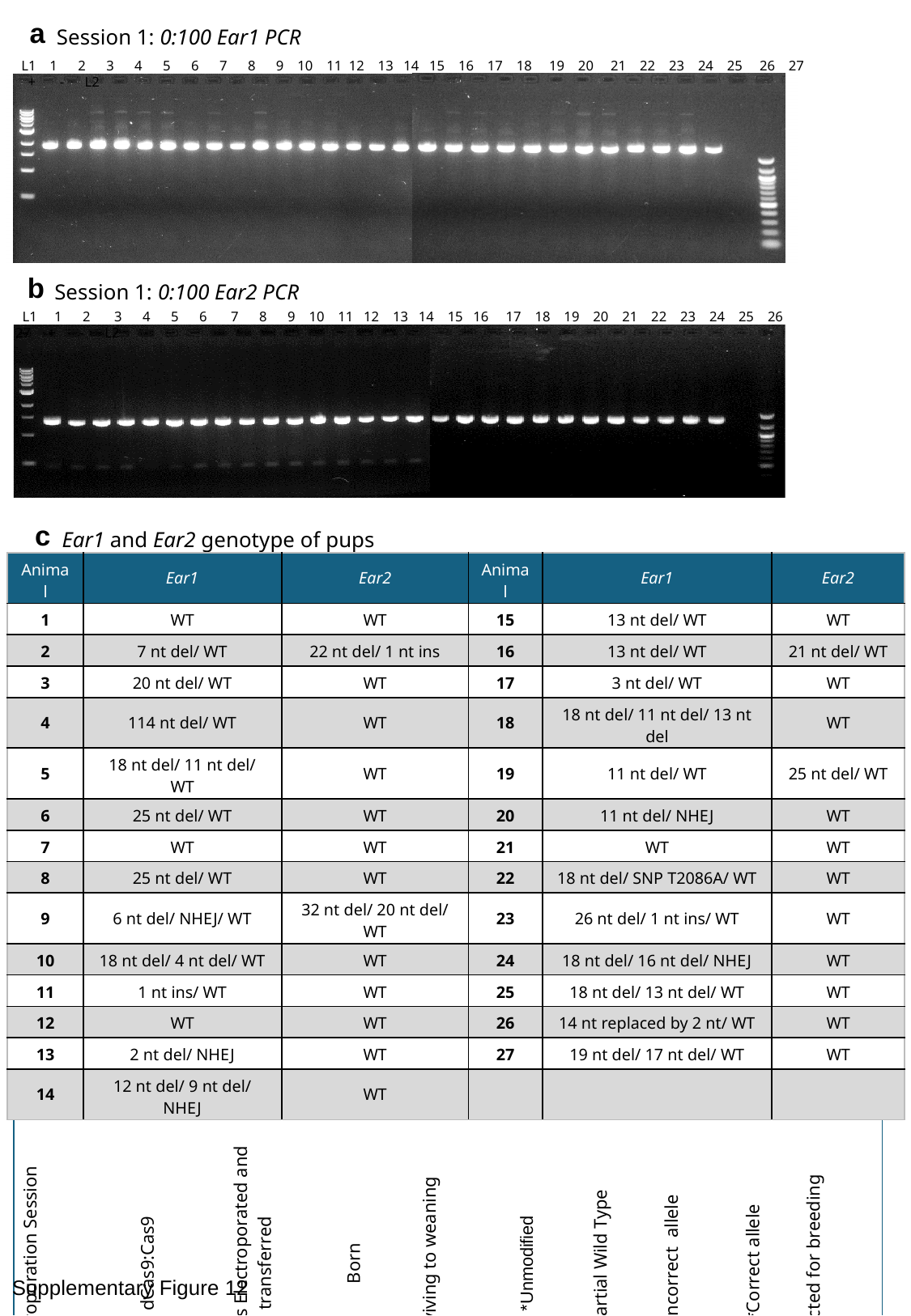

a
Session 1: 0:100 Ear1 PCR
 L1 1 2 3 4 5 6 7 8 9 10 11 12 13 14 15 16 17 18 19 20 21 22 23 24 25 26 27 + - L2
b
Session 1: 0:100 Ear2 PCR
 L1 1 2 3 4 5 6 7 8 9 10 11 12 13 14 15 16 17 18 19 20 21 22 23 24 25 26 27 + - L2
c
Ear1 and Ear2 genotype of pups
| Animal | Ear1 | Ear2 | Animal | Ear1 | Ear2 |
| --- | --- | --- | --- | --- | --- |
| 1 | WT | WT | 15 | 13 nt del/ WT | WT |
| 2 | 7 nt del/ WT | 22 nt del/ 1 nt ins | 16 | 13 nt del/ WT | 21 nt del/ WT |
| 3 | 20 nt del/ WT | WT | 17 | 3 nt del/ WT | WT |
| 4 | 114 nt del/ WT | WT | 18 | 18 nt del/ 11 nt del/ 13 nt del | WT |
| 5 | 18 nt del/ 11 nt del/ WT | WT | 19 | 11 nt del/ WT | 25 nt del/ WT |
| 6 | 25 nt del/ WT | WT | 20 | 11 nt del/ NHEJ | WT |
| 7 | WT | WT | 21 | WT | WT |
| 8 | 25 nt del/ WT | WT | 22 | 18 nt del/ SNP T2086A/ WT | WT |
| 9 | 6 nt del/ NHEJ/ WT | 32 nt del/ 20 nt del/ WT | 23 | 26 nt del/ 1 nt ins/ WT | WT |
| 10 | 18 nt del/ 4 nt del/ WT | WT | 24 | 18 nt del/ 16 nt del/ NHEJ | WT |
| 11 | 1 nt ins/ WT | WT | 25 | 18 nt del/ 13 nt del/ WT | WT |
| 12 | WT | WT | 26 | 14 nt replaced by 2 nt/ WT | WT |
| 13 | 2 nt del/ NHEJ | WT | 27 | 19 nt del/ 17 nt del/ WT | WT |
| 14 | 12 nt del/ 9 nt del/ NHEJ | WT | | | |
d
| | | | Pups | | | | | | |
| --- | --- | --- | --- | --- | --- | --- | --- | --- | --- |
| Electroporation Session | dCas9:Cas9 | Embryos Electroporated and transferred | Born | Surviving to weaning | \*Unmodified | \*Partial Wild Type | \*Incorrect allele | \*Correct allele | Selected for breeding |
| 1 | 0:100 | 114 | 29 | 27 | 4 | 18 | 4 | 19 | 3 |
Supplementary Figure 12

### Slide 20
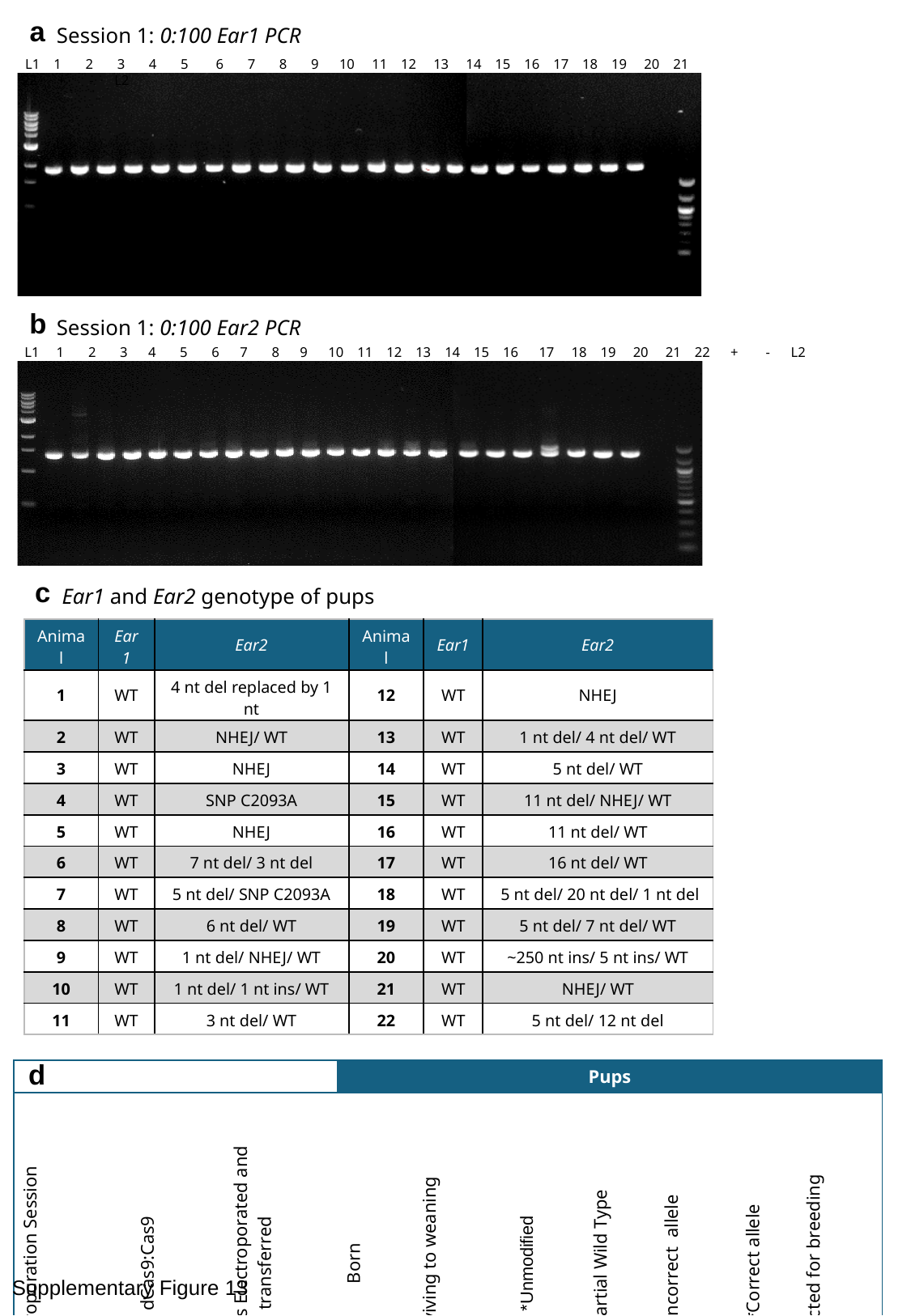

a
Session 1: 0:100 Ear1 PCR
 L1 1 2 3 4 5 6 7 8 9 10 11 12 13 14 15 16 17 18 19 20 21 22 + - L2
b
Session 1: 0:100 Ear2 PCR
 L1 1 2 3 4 5 6 7 8 9 10 11 12 13 14 15 16 17 18 19 20 21 22 + - L2
c
Ear1 and Ear2 genotype of pups
| Animal | Ear1 | Ear2 | Animal | Ear1 | Ear2 |
| --- | --- | --- | --- | --- | --- |
| 1 | WT | 4 nt del replaced by 1 nt | 12 | WT | NHEJ |
| 2 | WT | NHEJ/ WT | 13 | WT | 1 nt del/ 4 nt del/ WT |
| 3 | WT | NHEJ | 14 | WT | 5 nt del/ WT |
| 4 | WT | SNP C2093A | 15 | WT | 11 nt del/ NHEJ/ WT |
| 5 | WT | NHEJ | 16 | WT | 11 nt del/ WT |
| 6 | WT | 7 nt del/ 3 nt del | 17 | WT | 16 nt del/ WT |
| 7 | WT | 5 nt del/ SNP C2093A | 18 | WT | 5 nt del/ 20 nt del/ 1 nt del |
| 8 | WT | 6 nt del/ WT | 19 | WT | 5 nt del/ 7 nt del/ WT |
| 9 | WT | 1 nt del/ NHEJ/ WT | 20 | WT | ~250 nt ins/ 5 nt ins/ WT |
| 10 | WT | 1 nt del/ 1 nt ins/ WT | 21 | WT | NHEJ/ WT |
| 11 | WT | 3 nt del/ WT | 22 | WT | 5 nt del/ 12 nt del |
d
| | | | Pups | | | | | | |
| --- | --- | --- | --- | --- | --- | --- | --- | --- | --- |
| Electroporation Session | dCas9:Cas9 | Embryos Electroporated and transferred | Born | Surviving to weaning | \*Unmodified | \*Partial Wild Type | \*Incorrect allele | \*Correct allele | Selected for breeding |
| 1 | 0:100 | 120 | 22 | 22 | 0 | 13 | 9 | 13 | 2 |
Supplementary Figure 13
